# Improved inference and prediction of bacterial genotype-phenotype associations using pangenome-spanning regressions

**DOI:** 10.1101/852426

**Authors:** John A. Lees, T. Tien Mai, Marco Galardini, Nicole E. Wheeler, Jukka Corander

## Abstract

Discovery of influential genetic variants and prediction of phenotypes such as antibiotic resistance are becoming routine tasks in bacterial genomics. Genome-wide association study (GWAS) methods can be applied to study bacterial populations, with a particular emphasis on alignment-free approaches, which are necessitated by the more plastic nature of bacterial genomes. Here we advance bacterial GWAS by introducing a computationally scalable joint modeling framework, where genetic variants covering the entire pangenome are compactly represented by unitigs, and the model fitting is achieved using elastic net penalization. In contrast to current leading GWAS approaches, which test each genotype-phenotype association separately for each variant, our joint modelling approach is shown to lead to increased statistical power while maintaining control of the false positive rate. Our inference procedure also delivers an estimate of the narrow-sense heritability, which is gaining considerable interest in studies of bacteria. Using an extensive set of state-of-the-art bacterial population genomic datasets we demonstrate that our approach performs accurate phenotype prediction, comparable to popular machine learning methods, while retaining both interpretability and computational efficiency. We expect that these advances will pave the way for the next generation of high-powered association and prediction studies for an increasing number of bacterial species.

## INTRODUCTION

Bacterial genomics has recently entered an era of ‘big data’. Single cohorts with 10^4^-10^5^ samples, 10^8^ variants, and corresponding extensive high quality metadata are now publicly available (1, 2). In the context of bacterial populations, data inputs are typically genotypes or environmental factors (such as host age), and outputs are commonly antimicrobial resistance, host specificity or virulence phenotypes. With enough independent observations, these methods can automatically predict the phenotype of new isolates, and potentially tell us something about the underlying genetic mechanisms. A deluge of recent papers have applied general predictive models to such datasets, most showing high accuracy (3–8), though some commentaries have been more cautious in their conclusions (9, 10).

The overall problem of relating microbial genotype to phenotype has generally been approached by genome-wide association study (GWAS) methods (11–13). Generally these methods take a univariate approach, considering the association between a phenotype and a single variant, then ‘scanning’ along the whole genome one variant at a time. Microbial GWAS methods must incorporate a correction for population structure in each test, the specifics of which varies between methods and datasets. The simplicity of this method is one of its great strengths – it is quick and easy to apply, understand and visualise. Useful extensions which allow the estimation of heritability (14), the proportion of phenotypic variance explained by genotype, and prediction of phenotype (by forming linear predictors from significant variants, (15)) are also relatively simple to implement.

Large cohorts of bacterial sequences have been a tempting target for exciting new machine learning and ‘deep learning’ methods such as convolutional neural networks, which are able to relate arbitrary high-dimensional inputs to measured outputs with high accuracy, and without need for specialised model descriptions for each new problem (16). They are potentially broadly applicable to any problem with vast amounts of data, though perform best when the number of datapoints exceeds the number of dimensions. Unsurprisingly their uptake in sequence analysis (17, 18) and bacterial genomics specifically has been rapid (4, 5).

However, some issues crucial to understanding bacterial populations remain unaddressed. Firstly, bacterial populations tend to exhibit strong population structure, meaning samples cannot be treated as independent. In the context of prediction, this can result in the selection of features unrelated to the phenotype, but common to the background of associated strains (lineage effects). While not necessarily a problem in the training dataset, if new data is drawn from different strains this can lead to much poorer prediction than expected. This effect has been known since early efforts to statistically classify images, where from as early as 1964 it was shown that pictures of tanks were recognised by the background they appeared in, rather than the shape of the tank itself (19). In genetics, a similar problem has arisen due to an overrepresentation of samples of European ancestry in genotype databases. This has led to polygenic risk scores, which were originally thought to be highly accurate predictors of disease liability, to have significantly lower accuracy in non-European ancestry samples, which make up most of the global population (20). Additionally, these methods are unable to deal with missing input data. Naive variant calling in separate populations is likely to produce disparate sets of variants, and with very different minor allele frequencies. Without using a method which produces consistent variant calls in test datasets (not simply by cutting these out of a common call set), the accuracy of predictive models is likely to be heavily overstated.

Here, we set out to develop a method which combines the desirable attributes of both of these classes of approaches when analysing genetics underlying bacterial traits. We wished to retain the simplicity and interpretability of traditional univariate GWAS approaches, and combine this with the flexibility and accuracy of machine learning methods which can be fitted to the entire dataset at once. This multivariate approach reflects the polygenic nature of complex traits better than univariate methods. Unlike marginal tests, a multivariate regression approach gives rise to an increase of resolution when sample size increases, as has previously been noted in human GWAS (21–23). Additionally, simultaneously analyzing predictors together in a regression model means that interactions and correlation between the predictors (e.g. population structure) may be included implicitly.

Using large genomic datasets from four different species and sixteen varied phenotypes, we find that an elastic net model offers improvements over univariate GWAS, without sacrificing their major advantage of quantitative model interpretability. Using simulated data we demonstrate improved power and false-discovery rate at the single variant level compared with fixed and random effect models, and illustrate this use in practise on antibiotic resistance phenotypes in two species. We show further results which find similar accuracy between new machine-learning and simpler approaches, consistent with previous studies (4, 5, 24). Additionally, our approach was able to estimate trait heritability without assuming specific effect size distributions, which are unproven in bacterial populations. We argue that the prediction model itself is far less important than three other factors: the dataset itself, creating a method with careful genomic data management, and incorporating knowledge specific to bacterial populations.

Our approach models the entire pan-genome of the population, to include the large proportion of variation which resides in the accessory genome. It explicitly addresses issues of population structure, and consistent performance between trained and new (test) datasets. The method is broadly usable, not requiring programming knowledge or manual adjustments for new datasets, and allows for the sharing of models between researchers. We have implemented the association model in the pyseer microbial GWAS package, and consistent pan-genome variant calling in two further packages. An extensive tutorial for all of these methods is available online (https://pyseer.readthedocs.io/en/master/predict.html).

## MATERIALS AND METHODS

### Elastic net model

We use a high dimensional regression model which includes all genetic variants and covariates of interest 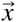. Typically this is not possible using classical inference methods such as the least squares estimator, as the number of genetic predictors *m* exceeds the number of samples *N*, leading to an under-constrained system. The elastic net defines such a function (25, 26), mixing L1 (lasso) and L2 (ridge) penalties, which is minimised with respect to the values of the intercept *b*_0_ and slopes 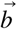:

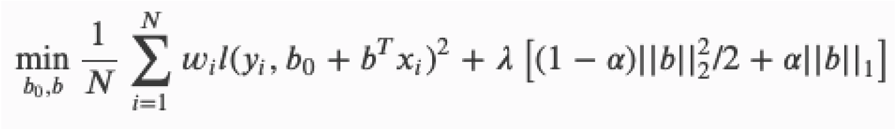

where *N* is the number of samples; *w*_i_ are positive weights (with a sum equal to *N*); *l*() is the link function (linear or logit respectively for continuous and binary phenotypes *y*_i_); λ > 0 is the magnitude of the penalty; 0 < α < 1 is the amount of mixing between L1 and L2 penalties. Minimising this function reduces the squared distance between predicted and observed values.

Given the strong linkage disequilibrium present in bacterial populations (11, 27), many genetic variants are strongly correlated across long distances, and it is therefore desirable to report all of this linkage block, rather than randomly selecting a representative from it. For this reason, the elastic net has been shown to be especially useful when the variables are dependent (25). We compare this selection with the lasso in our simulations. The value of α can be changed by the user, should they wish to opt for a more sparse model.

Two important parameters which are not directly set using the data are λ and α. The amount of penalty λ is set by cross-validation, using a default of 10 folds to pick the value of λ which maximises the *R*^2^ value of the model. The user can change the number of folds. The amount of mixing α between L1 penalties, which lead to sparse predictive models with mostly zero valued slopes, and L2 penalties, which lead to models with shrunken but non-zero predictors could also be chosen by cross-validation to prediction accuracy. However, in this application we propose using a value of around 0.01 throughout. A sparse model is preferred for prediction for speed and easier consistency between populations, but this leads to a loss of power in the context of GWAS, as potentially causally related predictors may be removed from the model (Figure 1, Supplementary figures 1-6).

**Figure 1:**
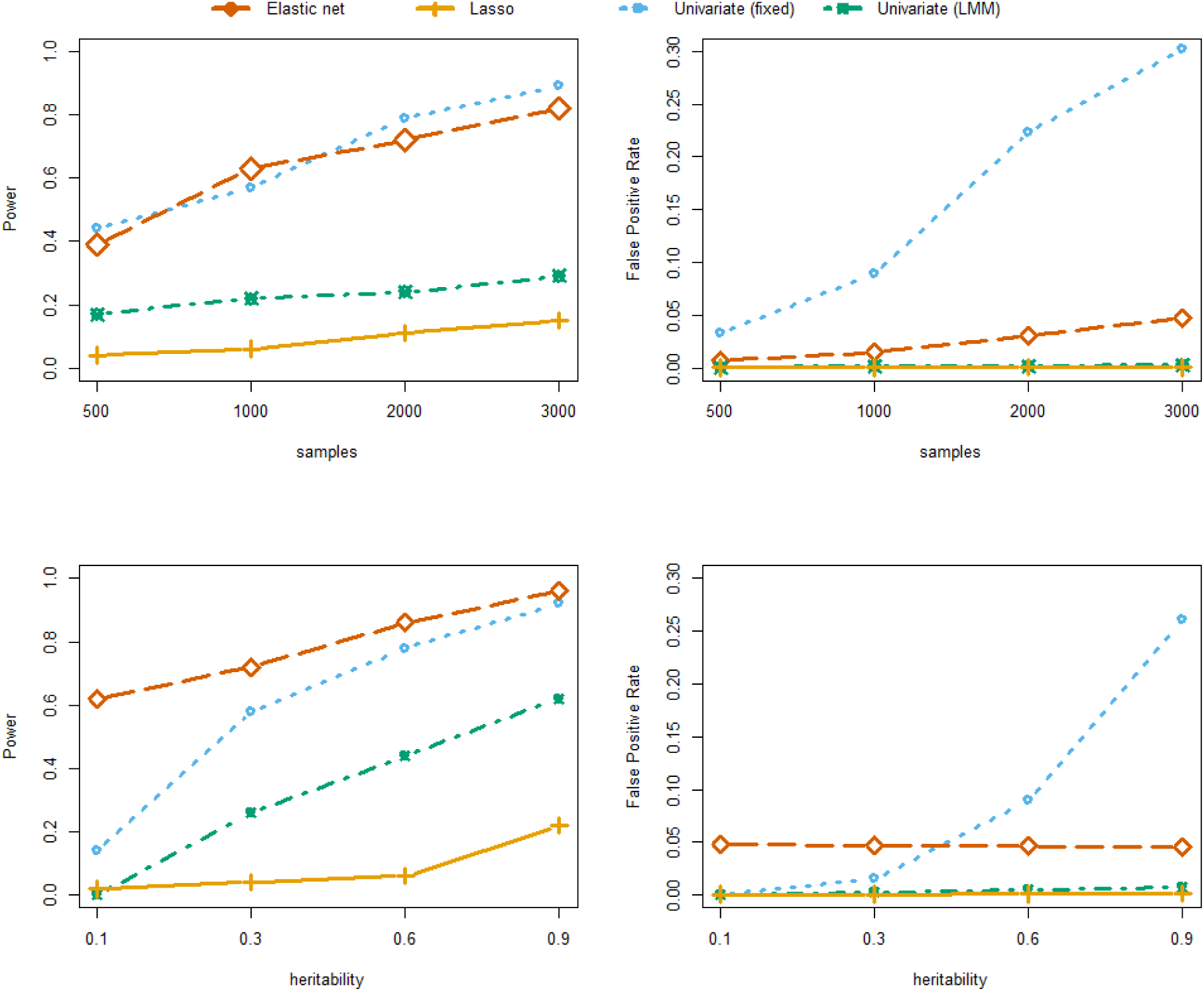
Power and false positive rates in the simulation study set up to resemble antibiotic resistance genotype-phenotype architecture. The top row shows the effect of sample size, with 100 causal variants in the *pbp2x* gene and a binary phenotype. The bottom row shows the effect of phenotype heritability, with 50 causal variants spread across the three *pbp* genes and a continuous phenotype. Multivariate methods tested were the elastic net with default ɑ (red diamonds), Lasso regression (orange lines). Univariate methods were the fixed effects/seer model (blue circles) and the FastLMM linear mixed model (green crosses).

By virtue of the fact that all pan-genomic variation enters this model, if a linear additive model of heritability is assumed, which it typically is in bacterial GWAS (28–30), the value of *R*^2^ calculated from the fit of the elastic net:

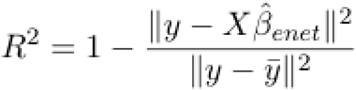

also serves as an estimate of the narrow-sense heritability *h*^2^. As *R*^2^ measures the variance explained by the model’s predictors, in this case all genetic features, this is equivalent to the proportion of phenotypic variance σ*_p_*^2^ explained by genetic variation σ*_g_*^2^, the definition of *h*^2^. This provides a lower-bound on *h*^2^ because the Lasso-type estimator is biased (31), and it tends to shrink some coefficients with weak effects towards zero, though these weak effects may have a significant effect on the trait variability.

### Efficiently modelling the entire pan-genome

Bacterial populations vary greatly in their sequence content, and mapping short variation within their core genes (coding sequences shared by all members of the population) is generally insufficient to capture all of the variation within the samples. In particular, accessory gene content has been shown to both vary independently of core variation (32), associated with clinically relevant phenotypes (12, 33), and be useful for predicting the evolution of the population (34, 35).

Early bacterial GWAS methods used k-mers, sequence words of fixed or variable length, to assay variation throughout the population independent of gene annotation or variant calling method (11, 12, 36). The set of common k-mers (1-99% frequency) is vast, particularly in the large and genetically diverse populations which are most amenable to GWAS. Efforts to model all of these sequence elements simultaneously are potentially computationally intractable, as these words will not fit in main memory, and model fitting takes an extremely long time.

We use two techniques to circumvent this issue while still including as much pan-genomic variation as possible. Following the idea of screening methods in ultra high dimensional data (37, 38), we use the absolute value of the sample correlation as a screening criteria for each variant:

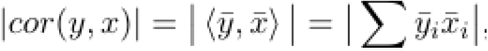

where *ȳ*, *x̄* are the standardizations of *y* and *x* such that

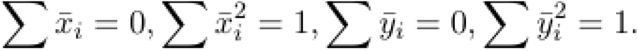

Using a single threshold on mean value for this correlation would lead to a large number of variants being removed before modelling, which is appealing computationally, but in our simulations led to a loss of power. We instead remove the lowest quartile, which maintained power, though did not reduce model size as much. The size of the quantile to remove can be set by the user.

We also follow the method used in DBGWAS (39), which after counting fixed-length k-mers constructs a de Bruijn graph of the population. Nodes in this graph are extensions of k-mers with the same population frequency vector, and whose sequence is referred to as unitigs. These unitigs greatly reduce the redundancy present in raw k-mer counts by combining those with the same patterns, and are generally easier to functionally interpret due to their longer length. We follow the same method as step 1 of the DBGWAS package, which uses the GATB library to construct a compressed de Bruijn graph (40), and then report frequency vectors of each unitig/node and unique pattern in a format readable by pyseer. We use a k-mer length of 31 throughout to count unitigs, as this was previously shown to maximise association power (39). This length can also be set by the user. We reimplemented this approach as a standalone package (https://github.com/johnlees/unitig-counter), also including tools to extend unitigs by traversing neighbouring nodes in the graph, and calculate distances between unitigs based on the graph using Dijkstra’s algorithm.

### Incorporating population structure

Population structure causes correlation between the genetic variants (unitigs) that make up 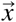. As all of these enter our model together, this effect may be implicitly controlled for without the need for further correction. We also wished to compare this to the use of an explicit correction term to test which approach is more effective. This can be included in the modelling step by a combination of three approaches: use of extra predictors in 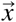 which account for population structure; modifying the per-sample weights *w*_i_; or by changing the folds used in cross-validation.

Fixed effect models typically use a principal components-type analysis to include main axes of variation in the population as covariates. In a new dataset, projection of variation onto these existing axes could be used, but would require large overlap between variant calls in each dataset to be accurate. Random effect models use a kinship matrix to include the sample covariance matrix in each association. For a new dataset, this would require calculation of covariance against the original dataset, which reduces model portability.

We therefore opted to use a definition of population structure which does not introduce extra predictors. This makes application of the model more straightforward in new datasets. Using a definition of clusters which naturally extends to new populations, which may have very different strain frequency and/or large numbers of novel clusters, further increases robustness in the face of between-dataset variation. Any method which produces discrete cluster membership definitions independent of cluster frequency is suitable for this purpose, such as sequence type, clonal complex or percentage identity cutoff. We opted to use the ‘strain’ definition provided by the PopPUNK software throughout our analysis due to its speed and biological basis (32), but our implementation allows any preferred definition of cluster membership to be used.

Given such a definition of cluster membership C(x), these clusters are then used as folds in the cross-validation step, which may be referred to as ‘leave-one-strain-out’ (LOSO) cross-validation. This is more realistic than random selection of folds, as random samples would maintain relative strain frequencies between training and test data, whereas new populations usually vary greatly in their genetic background (1, 10). Furthermore, we added the `--sequence-reweighting` option in pyseer, which defines the sample weights as being inversely proportional to the cluster/strain size:

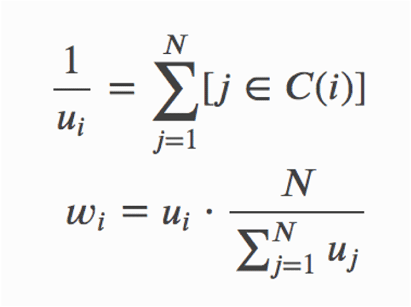

This ‘sequence reweighting’ is a commonly used definition in epistasis methods such as direct coupling analysis and correlation-based approaches, which have recently been successfully applied to genome-wide variation in bacterial populations (41–43).

### Phenotypic prediction while maximising consistency between datasets

Prediction of unobserved phenotypes *y*_i_ is achieved by forming a linear predictor of non-zero slopes in 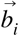:

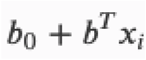

To which the appropriate link function *l*() is then applied to convert into a probability (which can be converted into a binary outcome using a threshold cutoff).

Predictors which are missing can either be ignored (set *x_i_* = 0) or imputed (set *x_i_* = *x̄*, where *x̄* is the allele frequency in the original dataset). For the unitig-caller approach described below, a missing call means genuine absence in the data, so we use the former approach. For variant calling methods where missing calls may be artifactual (such as SNP calls), we use the mean value imputation approach.

To apply fitted models to new populations, consistency in the construction of genomic variants 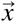 is important to maximise prediction accuracy. This can be highly challenging for short variant calls, but sequence element presence or absence is more amenable to this analysis. In general, de Bruijn graphs of different sample sets will have different node sequences, and therefore non-overlapping unitig calls. To solve this issue we instead check the test population for the presence of unitigs defined in the training population. To do this efficiently we create and save an FM-index for each input sequence (44), then search each these indexes in parallel for each unitig query (https://github.com/johnlees/unitig-caller). We implemented this in C++ using the SeqAn library (45).

For continuous phenotypes we report *R*^2^ to describe prediction accuracy. This is more difficult to interpret with binary phenotypes, especially in the presence of class imbalance and when the imbalance deviates from the population-wide prevalence. For binary phenotypes we found that reporting positive statistics such as sensitivity and specificity, or especially area under the curve (AUC), led to reports in the top decile for almost all datasets and methods, and were harder to intuitively compare between. We therefore report the false negative rate and false positive rate, along with the totals selected.

### Implementation details

We implemented the association model in version 1.3.2 of the pyseer bacterial GWAS package, which is written in python (46). This takes care of reading variants in many common formats, including the output from unitig-counter, as well as providing tools to help interpret associated sequences. We use python bindings to the fortran `glmnet` package to actually fit the model, as the use of warm-starts more efficiently solves the above equation at an array of values of λ (26). Cross-validation, parallelised if requested by the user, is used to select the value of λ with the greatest *R*^2^ value, as defined above.

Variants matrices are potentially very large, so to optimise speed and memory use we read these into a sparse matrix structure. Variants with allele frequency > 50% have their genotypes flipped to increase sparsity – these sites are flipped back during prediction. This sparse structure can be saved to disk to avoid repeated parsing of variant input files. Only haploid variant calls (0/1), allele frequency and sample order are saved in this file. After extracting the non-zero coefficients from the fitted elastic net, the input variant file is re-read with minimal parsing to output information about the selected variants. A SHA256 hash of the input file is calculated to ensure consistency with the original input file.

The fitted models are saved as an associative array, with variants names as keys (either sequence, or alleles combined with chromosome and position) and allele frequencies and fitted slopes as values. New variant call files are read with minimal parsing to extract just those sites which appear in the model, and at the end-of-file the appropriate imputation procedure is applied to model terms which were not found. This allows both rapid prediction in large new datasets, and an easy and portable way to share predictive models.

For fitted models, slopes, p-values (adjusted by any of pyseer’s other models) and allele frequencies are included in the output. Where the true phenotype is known, prediction accuracy is reported using *R*^2^, and a confusion matrix if the phenotype is binary. If clusters were provided, accuracy statistics for within each cluster are also included in the output.

The new code in pyseer includes automated tests and unit tests we wrote using test data distributed with the package. Documentation and a tutorial is available online (https://pyseer.readthedocs.io/en/master/predict.html). All three packages are available as source code and pre-packaged installation through conda.

### Preparation of datasets

Table 1 shows a summary of the datasets used in this paper. Sequence assemblies were available from the original publications, with the exception of one *N. gonorrhoeae* study (47). For this study we downloaded the read data, removed adapter sequences with trimmomatic (48) v0.36 and assembled with SPAdes v3.11 (49) using the `--only-assembler` and `--careful` options. For all datasets, we then called unitigs from each sample’s sequence assembly using a k-mer length of 31, and low frequency unitigs (AF < 1%) were discarded. For the Massachusetts and TB datasets additional genetic data was available. For the TB dataset, we used the variant call matrix provided by the study’s authors (5). For the Massachusetts and Maela datasets, we used SNP calls from an earlier GWAS in this population (50). Where a split into training and test data was needed, this was done at random in the ratio 2:1. When including a cluster assignment to account for population structure we used the previous assignments from PopPUNK where available (32). For TB we used the major lineage. When using a minimum inhibitory concentration for an antibiotic resistance, this phenotype was first log transformed before any downstream analysis. Other phenotypes were used as originally reported.

**Table 1:** Summary of datasets tested. Each dataset has a name it is referred to by in the rest of this paper. Most datasets have multiple phenotypes available, especially where multiple different antibiotic resistances are routinely phenotyped. Datasets without a training/test split were not evaluated for internal prediction ability, and all available samples were used to fit the model.

| Dataset name | Species | Phenotype(s) | Publication reference | Number of samples | Training/test | Number of genetic features |
| --- | --- | --- | --- | --- | --- | --- |
| TB | *Mycobacterium tuberculosis* | First line antibiotic resistance (rifampicin, isoniazid, pyrazinamide ethambutol) | (5) | 3566 | 2377 / 1189 | 6.4k (SNPs) |
| *N. gonorrhoeae* | *Neisseria gonorrhoeae* | Antibiotic resistance (azithromycin, cefixime, ciprofloxacin, penicillin, tetracycline) | (47, 51–53) | 1595 | NA | 550k (unitigs) |
| GAS | *Streptococcus pyogenes* | Virulence | (54) | 1730 | 1154 / 576 | 1.1M (unitigs) |
| SPARC | *Streptococcus pneumoniae* | Antibiotic resistance (penicillin, erythromycin) | (50, 55) | 603 | 400 / 203 | 90k (SNPs)<br>730k (unitigs)<br>10M (k-mers) |
| Maela | *Streptococcus pneumoniae* | Carriage duration<br>Antibiotic resistance (penicillin, erythromycin, trimethoprim) | (12, 28) | 3162 (antibiotic resistance)<br>2017 (carriage duration) | 1404 / 703 (carriage duration) | 121k (SNPs)<br>1.6M (unitigs) |
| GPS | *Streptococcus pneumoniae* | Antibiotic resistance (penicillin) | (1) | 5820 | NA | 1.7M (unitigs) |
| Netherlands | *Streptococcus pneumoniae* | Meningitis/carriage | (30) | 1837 | 1225 / 612 | 690k (unitigs) |

To generate simulated data used for testing power and false-discovery rate against a ground-truth, we simulated phenotypes but used observed SNP genotype data from the Maela dataset to ensure a realistic genetic model for bacterial population structure. Phenotypes were simulated using GCTA (14), either as continuous or as binary using a liability threshold model. Then, to assess the power of the methods with respect to the sample sizes, we randomly choose subsamples with 500, 1000, 2000 and 3000 samples.

## RESULTS

### Simulated phenotypes show advantages in power and false-discovery rate when using a multivariate approach

To test the characteristics of the elastic net compared to previous univariate GWAS approaches, we simulated phenotype data from the Maela population, using the observed genetic variation from this dataset. We chose either 5, 25, 100 or 300 true causal variants with the same effect size, either:

- Chosen uniformly at random across the genome, after LD-pruning (no variants with *R*^2^ > 0.9).
- Chosen uniformly at random from 1 to 3 pre-specified genes (*pbpX/pbp2x*, *penA/pbp2b*, *pbp1a*).

We chose the first setting to emulate a polygenic trait, with many variants of roughly equal effect associated across the genome. The second setting more closely resembles antibiotic resistance, where multiple alleles in either one or a small number of genes contribute to the effect. We ran the elastic net (α = 0.01), lasso regression (α = 1) as well as both univariate models (fixed effects and linear mixed model) previously implemented in pyseer. Variants were output by the genome-wide model if they have a non-zero coefficient, and in the univariate models if their p-value exceeds a significance threshold of 0.05 after Bonferroni correction. For each simulated dataset and method we calculated the power, the proportion of true causal variants in the output, and the false-positive rate, the number of variants selected in the output which are not true causal variants divided by the total number of variants being tested.

Firstly, our simulations were able to show that using the correlation filtering step (reducing input size by 25%) reduced power on average by 4% and 8% in the worst case (Supplementary tables 1-4), where many small effect variants are spread across the genome, and with no appreciable power loss with smaller or more concentrated causal variants. The sample correlation values are positively skewed due to population structure, so filtering all variants with sample correlation below the mean value rather than a quantile leads to an unacceptably high loss of power, as many causal variants would be removed.

Over all of our simulations, we found that either the elastic net or the fixed effect univariate model had the highest power depending on the setting, and both always had higher power than the linear mixed model (Figure 1 and Supplementary figures 1-6). The elastic net performed better in situations where the heritability was low, or causal variants were spread out across the genome. There was slightly lower power for all methods with binary phenotypes, and this increase was more pronounced in the linear mixed model, possibly due to being the only model that used a Gaussian error structure in both settings.

However, in exchange for reduced power, the linear mixed model consistently showed the best control of false positives in all settings, always below 5%. In contrast the fixed effect model had a much greater false-positive rate than any other method, which grew both with sample size and heritability. The elastic net’s false-positive rate was typically below 5% and was robust across the ranges of heritability tested, though did increase slightly with larger sample sizes as more variants were included in the fitted model. With such a large number of variants even a small false-positive rate can be problematic, so combining with a ranking by p-value is important. It is possible to do this with least angle regression (56) in lasso regression, but in this case it is simpler to provide this ranking from one of the univariate tests.

### Whole-genome model of bacterial phenotypes enables heritability estimation and clearer association mapping

We also tested lasso regression as a GWAS method. For smaller numbers of causal variants performance was similar to the linear mixed model both in terms of power and false-positive rate, in some cases having slightly higher power. However when the number of causal variants was higher the amount of sparsity introduced was too high, reducing power below other methods (though false-positive rate was low in all settings for the same reason). As the number of causal variants is generally not known a priori, we would therefore always recommend the elastic net over the lasso.

These results show that the elastic net can be used as an effective tradeoff between the regimes of the two commonly used univariate models, having higher power than the linear mixed model and a lower false positive rate. As many bacterial GWAS results must be followed up with lab work, these results suggest a dual-approach of variable selection with the elastic net followed by ranking results with the linear mixed model may be useful. This is possible in a single step in pyseer.

#### GWAS of pneumococcal antimicrobial resistances

Next, we tested our method for GWAS on phenotypes with a known cause, and compared our results to those from previous univariate approaches. First, we analysed a well studied phenotype and dataset: sensitivity/resistance to β-lactams in the SPARC cohort of 603 *S. pneumoniae* genomes. Resistance is mainly conferred by allelic variation of three genes (*pbp1a*, *pbp2b*, *pbp2x*) which are easily detected by most GWAS and machine-learning methods with SNP calls as input (50, 57), though the specific variants detected are not identical between methods (58).

Figure 2 shows the results of this analysis. With both methods *penA*/*pbp2b* and *pbpX*/*pbp2x* are clearly the strongest hits. *pbp1a* is also selected by both methods, and while it can be seen on both Manhattan plots it is slightly clearer in the gene summary plot for the elastic net, due to the larger number of SNPs selected in the gene. Taking hits with a p-value above a threshold results in a very clean result for the LMM with this strongly selected phenotype, only a few non-causal genes are included, usually with only a few SNPs and much lower ranked than the causal genes. The elastic net selects many more non-causal SNPs across the MAF spectrum, though combining with p-values and number of hits allows these to be effectively filtered. It should be noted that effect size does not appear to be an effective filtering criterion without taking into account minor allele frequency, which may have implications for other machine-learning methods where a p-value cannot easily be integrated. Both methods calculate a comparable heritability estimates for this trait, for the LMM *h*^2^ = 0.89, and for the elastic net *h*^2^ = 0.81.

**Figure 2:**
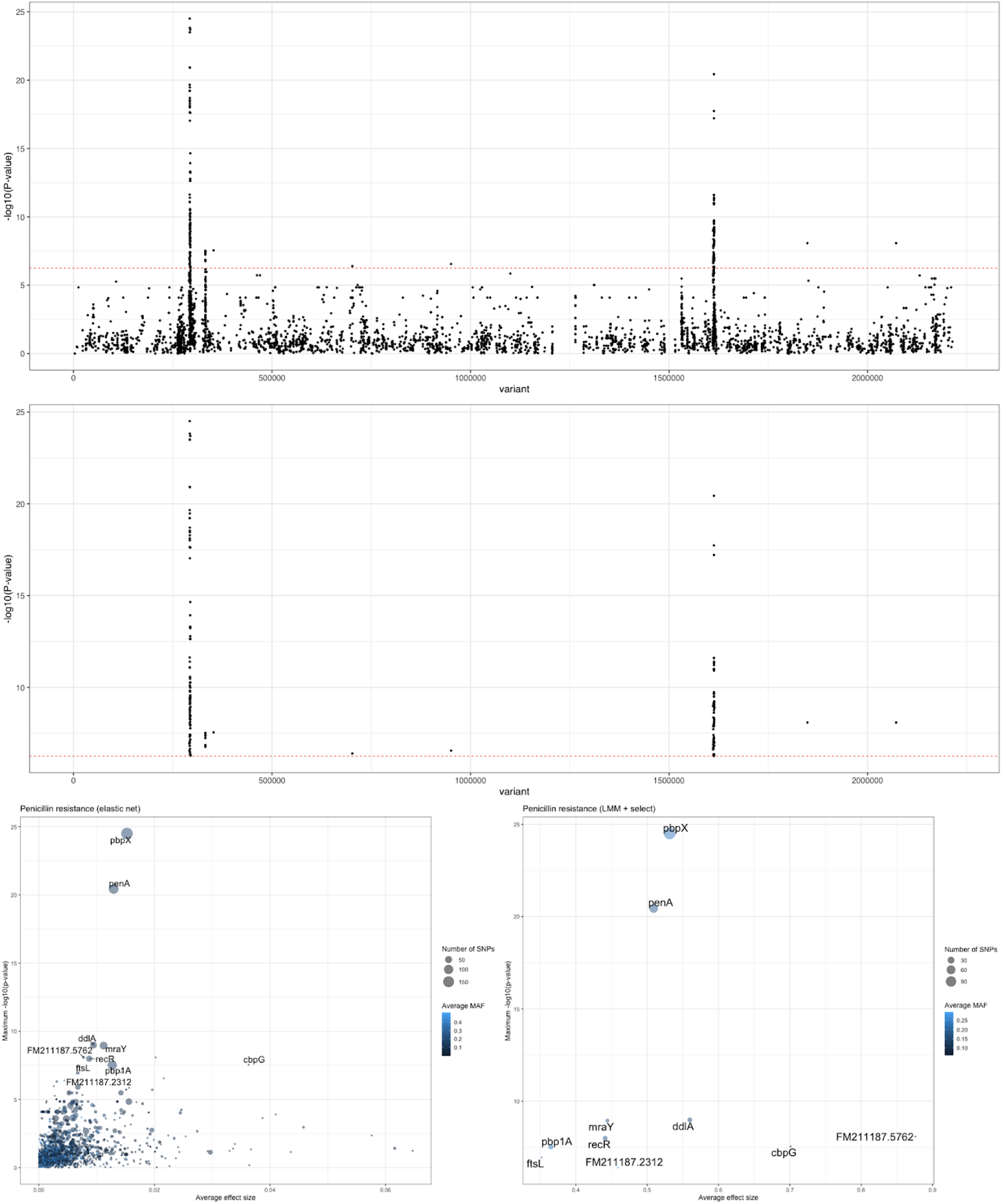
Elastic net and linear mixed model with SNP-based penicillin resistance. The top panel shows a Manhattan plot of the selected elastic net variants, with a Bonferroni-adjusted significance threshold in red. The three biggest peaks are in the causal *pbp* genes. The middle panel shows the same result with the univariate LMM, taking only those SNPs above the significance threshold. The bottom panels show a summary of the genes selected by both methods (left – elastic net; right – LMM), averaging p-value and effect size within each gene.

We then undertook a more challenging analysis, using unitigs to investigate antimicrobial resistances in the larger Maela cohort (3162 *S. pneumoniae* genomes). We have previously attempted GWAS on this dataset using the fixed effects model, in the original description of our SEER software (12). We did not run on tetracycline resistance or chloramphenicol resistance, as these are driven by single elements which were easily detected with the previous method. We instead used trimethoprim and erythromycin resistances. Trimethoprim resistance is expected to have two causal loci (*folA/dyr*, *folP*) due to being administered jointly with sulphamethoxazole. Indeed, using our method on trimethoprim, the two causal loci are clearly identified and are the most highly ranked on a Manhattan plot (Supplementary figures 7 & 8); applying sequence reweighting makes little difference to the result. Erythromycin resistance has multiple causal mechanisms (*ermB*, *mel*, *mef*) which were not easily found in our previous attempt. Again, the erythromycin results contained many peaks in their Manhattan plots (Supplementary figures 9 & 10). Mapping the unitigs directly to resistance genes did find significant results in *ermB* (9 hits, min p = 10^−47^), *mel* (12 hits, min p = 5×10^−42^) and *mef* (6 hits, min p = 5×10^−42^), and while this was clearly more successful than our previous analysis, when considering the noise when mapping to a single reference these causal mechanisms would not stand out (Supplementary figures 11 & 12). So while our method reduced the computational burden of the analysis, it was not able to easily resolve the causal mechanisms in this challenging example. This suggests that both univariate tests and black-box machine learning type approaches would struggle to arrive at true causal predictions under such circumstances.

#### Heritability and mapping of gonococcal resistance

We also applied our method to a combined cohort of 1595 *N. gonorrhoeae* genomes where resistance to five different antibiotics has been measured. This data has previously been used to do GWAS using a LMM, with selected loci then entering a reduced dimension epistasis analysis (53).

The mapping of resistance genes for these antibiotics using our approach was also successful. The original analysis looked at ~8700 SNPs with MAF > 0.5%; we used 5.3×10^5^ unitigs with MAF > 1%. Azithromycin (AZI) had 4612 unitigs selected, with the top hits mapping to the four 23S rRNA sequences in the genome. The original analysis only identified a SNP in one of these repeated rRNA sequences, likely due to the impossibility of mapping variation in these repeats at a single base level – this is an advantage of being able to report multiple mappings of sequences at the final stage. Cefixime (CFX) identified the *penA* region, as in the original analysis, and also suggested an association in the promoter of *opaD*. Ciprofloxacin (CIPRO) had hits throughout the genome, as in the original analysis and similar to the analysis of erythromycin above – combining the LMM with the elastic net may reduce candidate regions in these cases. Penicillin (PEN) had a hit in the *porB* region, as in the original analysis, along with hits in *lgtE*, *mexB* (and efflux pump) and a prophage. Tetracycline (TET) similarly had a replicated hit in the *porB* region, along with the *cysN* promoter, an alternative *pilE* allele and *remE* (a ribosomal methyltransferase). Our method therefore broadly replicated the results from the LMM, and added new candidate hits, as expected.

We also calculated the narrow sense heritability *h*^2^ using our elastic net and unitig method, and compared these to previous methods (figure 3). Our estimates were very similar to those of *h*^2^_SNP_ from the original paper, though consistently slightly higher (3% ± 7%), which may be a result of including more of the population variation through unitigs. Using a simple estimate of shared sequence content as the kinship matrix led to a likely overestimate of heritability.

**Figure 3:**
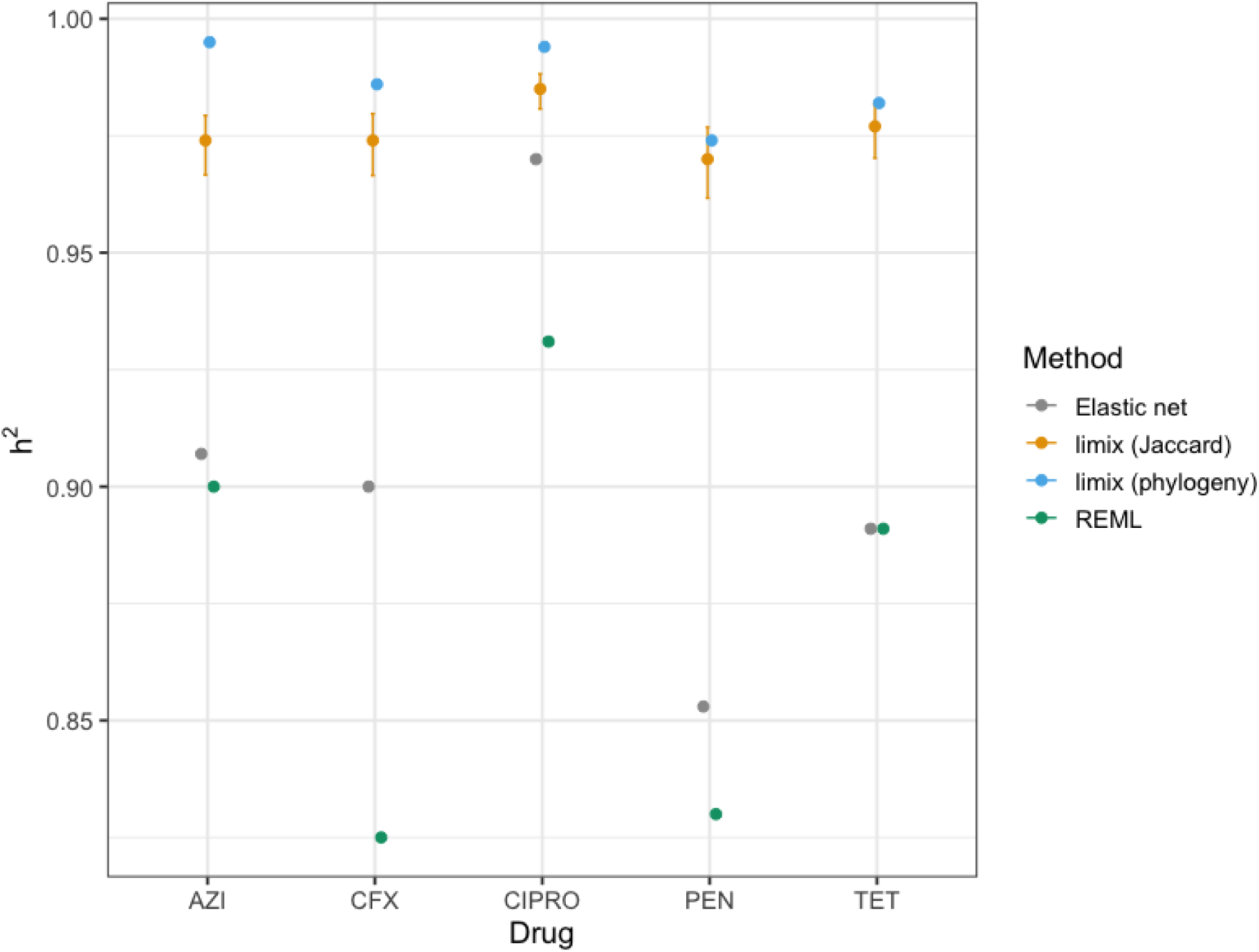
Heritability estimates for antibiotic resistance in the combined *N. gonorrhoeae* dataset, using different methods. For each of the five antibiotic resistances measured in this dataset we report the narrow-sense heritability (*h*^2^) from our elastic net method and unitigs (grey), the limix method implemented in pyseer, using sequence distance (gold) or phylogeny (blue), and the restricted maximum likelihood (REML) approach used in the original publication (green). For limix estimates, 95% CIs were calculated with FIESTA (59). These are not shown for the phylogeny method as they span a range wider than the plot (0.11-1).

### Accurate prediction within and between cohorts without sacrificing model interpretability

These models, shown above to be easily interpretable using standard methods from bacterial GWAS, can also be used to construct a linear model to predict phenotypes in new data. In this section we evaluate these predictions compared to other models and variant calling methods using a variety of datasets and phenotypes.

#### M. tuberculosis resistance is equally well predicted by the elastic net

We first evaluated the predictive performance of these models compared to more complex models, with and without population structure control. We used an *M. tb* dataset with antibiotic resistance to four first-line antibiotics (rifampicin, isoniazid, ethambutol and pyrazinamide). As *M. tb* has no accessory genome and minimal core gene variation (60), comparison with more complex models and a SNP alignment is possible. Previous work has evaluated the use of a multitask deep neural network, and when comparing this to lasso regression found comparable accuracy (5). Using the same input of ~6500 short variants across the allele-frequency spectrum for these 3566 samples (split into training and test datasets) led to an average false-negative rate of 2% ± 3% in the unweighted model 3% ± 4% in the weighted model, and false-positive rate of 11% ± 8% in the unweighted model, 12% ± 10% in the weighted model (Supplementary table 5). The elastic net therefore gives similar performance to the lasso as shown previously, as well as the more complex neural network. It is however much easier and faster to run on standard hardware, and gives GWAS results which are far more readily interpretable.

In this case, the weighting generally performs slightly worse than an unweighted model. The population structure of this sample is relatively simple, with four distinct lineages. This is likely well-captured implicitly by the unweighted model, so the categorical weighting is of lower resolution. Sequence reweighting is instead expected to be more effective in datasets with more complex structures, or when adding in samples which are genetically distant from the training set (42, 43), which we explore this further in the next section. Applying these weights did allow us to easily see that the majority of errors occurred in lineage I, which has deep branches forming genetically separated subclades, with generally perfect prediction in the other three lineages.

#### Prediction of pneumococcal resistance using different variant types

We also investigated the advantages of the use of unitigs over other variant calling methods. Using the same SPARC dataset described above for β-lactam and erythromycin resistance, we compared computational resources and prediction accuracy using SNPs, k-mers and unitigs (table 2).

**Table 2:** Predicting antibiotic resistance in the SPARC collection using different variant types. Using a training:test split of 2:1, prediction accuracy of two phenotypes was tested using 90k SNP calls from mapping to a reference genome, and with 730k unitigs. We also tested prediction using 10M variable length k-mers to illustrate the heavy computational resource use in even a relatively small dataset.

| Variant type | Phenotype | Number selected | False-positive rate | False-negative rate | CPU time | Memory usage |
| --- | --- | --- | --- | --- | --- | --- |
| SNPs | β-lactam | 4374 | 3% | 7% | 4.4 mins | 1.3 Gb |
|  | Erythromycin | 2341 | 3% | 63% | 4.1 mins | 1.3 Gb |
| Unitigs | β-lactam | 8247 | 5% | 7% | 49.7 mins | 18 Gb |
|  | Erythromycin | 1591 | 9% | 39% | 52.6 mins | 6.9 Gb |
| K-mers | β-lactam | 15121 | 6% | 7% | 7.0 hrs | 212 Gb |

We found that for β-lactam resistance all three variant types gave similar predictive accuracy, with the elastic net able to select a small proportion of the total input variants in each case, and apparently fairly insensitive to the far greater noise present in the higher dimensional variant types. As this resistance is due to allelic variation in core genes we expect all three types to tag the causal variation equally well. For erythromycin, where causal variants are not all found in core genes, we observed a reduction in false negative rate when using unitigs. Computational usage increased roughly as *NM* (*N* number of samples; *M* number of variants). For common variants, *M* reaches an asymptote for a given population, the main requirement is therefore based on *N*. For all methods CPU time was modest, but memory usage may pose a problem. SNPs are tractable on a laptop, but unitig analysis likely requires a compute cluster for the model fitting (using a fitted model on test data requires negligible resources). K-mers require an enormous amount of memory, which would not scale to larger datasets. Though the unitig analysis was easy to schedule on our cluster, future improvements to reduce memory use could include accessing the variants as they are needed from disk, or fitting the elastic net in chunks (61).

#### Reduced inter-cohort accuracy is ameliorated with consistent genetic calls and population structure control

Random splits of single datasets in test and training data, while convenient for analysis, may mask inter-dataset differences such as class imbalance (different resistance rates), unobserved lineages, and technical errors (variant calling) (10). To test a more realistic example, where a previously fitted model is used to predict resistance status in new, unobserved data, we set up a prediction experiment using genomic data from three large, very different pneumococcal cohorts with β-lactam resistance: SPARC (603 US children covering introduction of vaccine); Maela (3162 unvaccinated infants and mothers); GPS (5820 globally distributed samples, mostly vaccinated). We counted unitigs for each population and used these to train a predictive model. These models were evaluated on the data they were trained on, and on the other two cohorts by using consistently named unitigs from unitig-caller (table 3).

**Table 3:** Comparison of intra- and inter-cohort prediction accuracy. For each prediction the error rates are listed along with overall R^2^. Cells in green are within-cohort. The first three rows show results with sequence reweighting; the second three rows without weighting. For SPARC and Maela phenotype was binary (resistant/sensitive), for GPS phenotype was continuous (MIC). Where conversion was needed, we applied the standard breakpoint of MIC > 0.12 mg/L for resistance. FNR – false-negative rate; FPR – false-positive rate.

| Model | Number of selected unitigs (% in pbp genes) | SPARC data |  | Maela data |  | GPS data |  |
| --- | --- | --- | --- | --- | --- | --- | --- |
| SPARC (sequence reweighting) | 5251 (10%) | FNR | 0.063 | FNR | 0.007 | FNR | 0.149 |
|  |  | FPR | 0.024 | FPR | 0.239 | FPR | 0.134 |
| | | $R^2$ | 0.837 | $R^2$ | 0.439 | $R^2$ | 0.505 |
| Maela (sequence reweighting) | 6645 (14%) | FNR | 0.446 | FNR | 0.082 | FNR | 0.029 |
|  |  | FPR | 0.005 | FPR | 0.042 | FPR | 0.382 |
| | | $R^2$ | 0.276 | $R^2$ | 0.760 | $R^2$ | 0.425 |
| GPS (sequence reweighting) | 894 (4%) | FNR | 0.011 | FNR | 0.144 | FNR | 0.094 |
|  |  | FPR | 0.411 | FPR | 0.177 | FPR | 0.200 |
| | | $R^2$ | 0.447 | $R^2$ | 0.458 | $R^2$ | 0.545 |
| SPARC | 7261 (10%) | FNR | 0.040 | FNR | 0.012 | FNR | 0.165 |
|  |  | FPR | 0.013 | FPR | 0.163 | FPR | 0.130 |
| | | $R^2$ | 0.901 | $R^2$ | 0.487 | $R^2$ | 0.487 |
| Maela | 8705 (9%) | FNR | 0.397 | FNR | 0.063 | FNR | 0.049 |
|  |  | FPR | 0.011 | FPR | 0.036 | FPR | 0.322 |
| | | $R^2$ | 0.339 | $R^2$ | 0.805 | $R^2$ | 0.449 |
| GPS | 7511 (2%) | FNR | 0.050 | FNR | 0.319 | FNR | 0.129 |
|  |  | FPR | 0.152 | FPR | 0.026 | FPR | 0.037 |
| | | $R^2$ | 0.656 | $R^2$ | 0.452 | $R^2$ | 0.864 |

Between-cohort predictive accuracy was considerably lower than within-cohort accuracy, but still outperforms an intercept only model in all cases. The use of unitigs proved successful – repeating the SPARC-Maela comparisons with SNPs led to sensitive predictions for every sample, as the selected SNPs were called as missing in the other cohort. To fix this issue with SNPs would likely require a labour intensive mapping and joint recalling of variation, whereas using sequence elements can use the simple search implemented in unitig-caller. Depending on the specific model and dataset combination, errors can be much more commonly type I or type II, possibly reflecting class imbalance, despite overall resistance rates in the pneumococcus being stable (62). The GPS cohort gave the worst performing model, despite it being the largest collection. This is a very genetically diverse sample, which introduces more potential for confounding lineage effects to enter the model.

Furthermore, this cohort is a mix of sequences isolated from both asymptomatic carriage and disease, whereas SPARC and Maela contain only asymptomatic carriage. The GPS cohort is enriched with more virulent strains, which have more frequently faced treatment with antibiotics, and have a higher rate of resistance (1, 63).

We also note that the AUC of the ROC curve is misleadingly high (0.9185/0.9728 for the weighted/unweighted GPS model), and would encourage the reporting of error rates as more intuitive summaries of accuracy for bacterial traits such as resistance.

We found that sequence reweighting generally reduced prediction accuracy for this phenotype, although the LOSO strategy within-cohort gave more representative accuracy estimates for out-of-cohort prediction, and more of the selected variants were in the causal loci. There was little difference in accuracy between using mean AF imputation for missing unitigs, versus calls of absence for all samples.

#### Virulence phenotypes can be predicted with sequence reweighting preventing overestimation of accuracy

Most work on prediction of bacterial phenotypes have focused on antibiotic resistance, but many more complex phenotypes relating to bacterial virulence are now available. For these phenotypes, which are under weaker or no selection, instead of a few strong effects, multiple smaller effects are expected in the genome (28, 30, 54). Therefore, a model which may include more of these effects, which would be missed with a p-value threshold, may be expected to perform well.

We applied our method to predict the duration of asymptomatic carriage in a subset of the Maela cohort, which can easily be visualised in the manner of a linear regression (figure 4). We show the observed versus predicted values for the training and test sets, both with and without sequence reweighting. In the unweighted test set R^2^ (and *h*^2^) was 0.89 (top left panel), but the test R^2^ was only 0.27 (bottom left panel), showing clear overfitting. With sequence reweighting and LOSO, the training and test estimates were much closer (right panels, 0.37 and 0.28, respectively). In this case, sequence reweighting gave a more realistic heritability estimate. *h*^2^ was previously estimated to be 0.634 using phylogenetic pairs, and 0.445 using REML – these may be overestimates, especially as these revised estimates used information from more of the genome.

**Figure 4:**
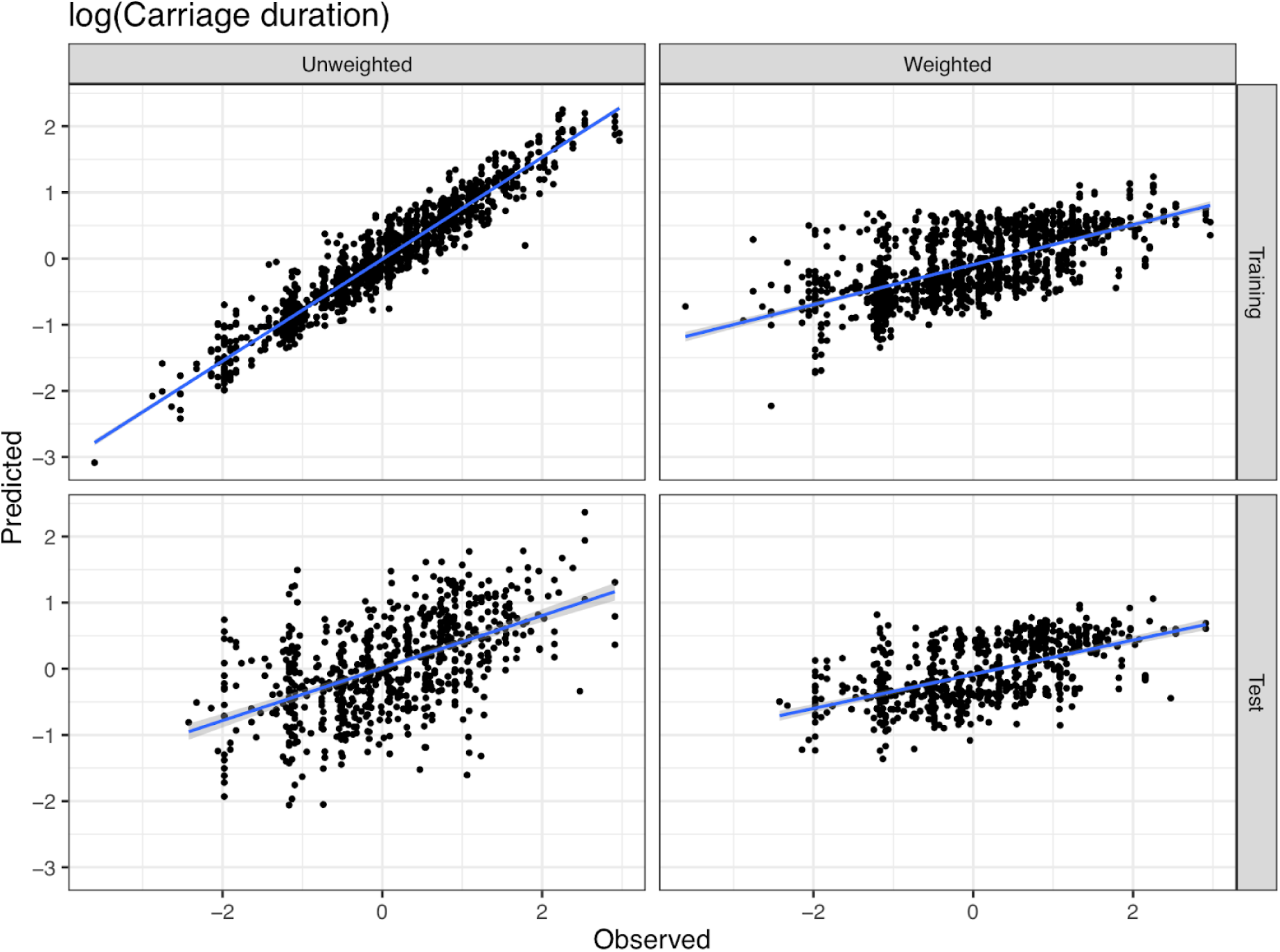
Prediction of carriage duration and the effect of sequence reweighting on heritability estimation. For the same training:test split, each panel shows observed log(carriage duration) values on the x-axis, and model-predicted values on the y-axis, with a fitted linear regression. The left column shows the unweighted model on the training data (top) and test (bottom); the right column shows the same for the model with sequence reweighting.

We also tested virulence prediction in two *Streptococcal* species, which would be a useful application for routine pathogen surveillance, but has not been as thoroughly explored as resistance prediction. Using *S. pneumoniae* isolated from Dutch adults, we fitted a model which selected 9701 unitigs. This model was able to predict meningitis versus carriage genomes (test FPR: 0.059; FNR 0.12) and gave a similar *h*^2^ estimate to that originally reported (0.65 vs. 0.70). Comparing tissue infection with carriage of *S. pyogenes* gave a model with 5817 unitigs which had a higher error rate than the *S. pneumoniae* model (test FPR 0.24; FNR 0.25), but the phenotype had a correspondingly lower *h*^2^ estimate of 0.343.

## DISCUSSION

In this paper we developed a microbe-specific implementation of a machine-learning tool, and showed how this can be used to better understand the link between bacterial genetic variation and phenotypic variation. Along with the statistical advantage of introducing a full-genome multivariate model, we further addressed three key aspects of these datasets while producing our open-access software tool. Pangenomic variation was covered using a unitig definition of population variation, which we showed to be scalable, unlike k-mers, and better suited to analysing accessory genome variation and inter-dataset consistency, unlike SNPs. Population structure was accounted for using the implicit properties of the elastic net, as well as explicit options of sequence reweighting and leave-one-strain-out cross-validation. We maintained a clear link between our resulting models and underlying genetics by combining linear models with a suite of tools to interpret the variants selected in the model. This had the further advantage that it significantly reduced the number of sequence elements to be processed after association. Using selected unitigs allowed for a much smoother use of the interactive plotting software phandango (64), which is one of the fastest ways to interpret bacterial GWAS results.

Our method can be used in a GWAS context, as well as for prediction. We showed superior performance to univariate models in simulated settings, and using real data from two species showed how this method can be combined with previous approaches to understand resistance and epistasis. We also obtain useful estimates of trait heritability, some of which show evidence that previous approaches may have overestimated this quantity. For the purposes of prediction, on a simple dataset we replicated the result that regularised linear models perform similarly to more flexible deep learning methods. We also applied our method to a range of datasets from different species and phenotypes, including resistance, carriage duration and virulence. Though our models generally performed well when measured on error rates, an experiment with models on three separate cohorts showed how accuracy falls outside of the target dataset. External datasets may have different strain compositions due to different biases toward more or less virulent strains, geographical separation, vaccine use or antibiotic consumption in the population. We would reiterate the caveat that while these models can be useful, high accuracy on test data should not be taken as a general measure of confidence (9). Batch differences such as genotyping methods between cohorts exaggerate this problem, so a consistent approach (such as the one we provide here) should be used. Unsurprisingly, curated resistance sets – the result of decades of research – still generally perform better, although even this in silico method loses accuracy between datasets (24). Less well understood and potentially polygenic phenotypes such as virulence offer an attractive target for our model, as we demonstrated on two Streptococcal pathogens.

Along with these theoretical advances, our package has a number of practical advantages. All of the elements of our method are freely available, well-documented and part of a continuous unit-testing framework. Users can construct and evaluate models easily, without the need for programming experience, with options which retain the flexibility to modify the model parameters. There is no need for specialist hardware such as the graphics cards needed to fit large deep learning models. The models themselves are saved in a human-readable format, and are easy to share and reuse and have minimal hardware-specific requirements.

More broadly, we have considered techniques which are routinely used in the analysis of modern datasets, which are generated continuously with high-throughput methods. These can be adapted to perform fundamental tasks in bacterial genomics in ways which are useful, and that scale with our ambitions to discovering causal drivers and predicting phenotypes from the genome variation. Collections of high quality whole genome sequences are now available at a scale that would have been unfathomable just a few years ago. Many of these datasets are publicly available already, and many more are being generated from new, larger projects and routine surveillance by public health agencies. Care must be taken to assure the unique properties of bacterial populations are properly modelled, and that we use appropriate measures of success. Complex models should be compared to simple models (65) – not just in terms of accuracy, but also for their ability to look at underlying biology. In many cases the limiting factor is unlikely to be model flexibility. With our pangenome-spanning penalized regression models, we hope to have made a useful and usable contributions that respect these principles.

## AVAILABILITY

The pyseer package is available as source code at the GitHub repository (https://github.com/mgalardini/pyseer; Apache 2.0 license), documented on readthedocs (http://pyseer.readthedocs.io/), and available for install on bioconda (https://anaconda.org/bioconda/pyseer). The unitig-counter package is available at the GitHub repository (https://github.com/johnlees/unitig-counter; AGPL 3.0 license), and available for install on bioconda (https://anaconda.org/bioconda/unitig-counter). The unitig-caller package is available at the GitHub repository (https://github.com/johnlees/unitig-caller; Apache 2.0 license), and available for install on bioconda (https://anaconda.org/bioconda/unitig-caller).

## ACKNOWLEDGEMENTS

We thank the developers of DBGWAS who wrote the majority of the code to count unitigs, which we modified for this application. We are particularly grateful to Leandro Ishi Soares de Lima and Rayan Chiki for helping test and debug code from DBGWAS and GATB used in unitig-counter. We also wish to thank the developers of SeqAn 3.0 for their help implementing the FM-index in the unitig-caller package. We would like to thank Benjamin Schubert and Deborah Marks for providing the phylogeny for the *N. gonorrhoeae* dataset. Daniel Falush provided useful feedback on an earlier presentation of this work.

