## Supplementary information for "Improved inference and prediction of bacterial genotype-phenotype associations using pangenome-spanning regressions"

#### Contents

**1 Supplementary tables** **2**

**2 Supplementary figures** **7**

### 1 Supplementary tables

| | thresholds | 500 | 1000 | 2000 | 3000 |
|---|---|---|---|---|---|
| 5 snps | the 1st Qu. | 5 (90248) | 5 (90746) | 5 (90658) | 5 (90742) |
| | the median. | 5 (60411) | 5 (60506) | 5 (60507) | 5 (60507) |
| | the 3rd Qu. | 4 (30247) | 5 (30252) | 4 (30254) | 5 (30254) |
| 25 snps | the 1st Qu. | 25 (90690) | 25 (90425) | 25 (90721) | 25 (90758) |
| | the median. | 23 (60216) | 20 (60506) | 23 (60507) | 22 (60501) |
| | the 3rd Qu. | 14 (30222) | 15 (30254) | 15 (30254) | 15 (30254) |
| 100 snps | the 1st Qu. | 94 (90273) | 95 (90733) | 94 (90739) | 95 (90729) |
| | the median. | 74 (57564) | 77 (60487) | 79 (60506) | 74 (60427) |
| | the 3rd Qu. | 49 (30225) | 50 (30254) | 51 (30254) | 48 (30254) |
| 300 snps | the 1st Qu. | 273 (87892) | 284 (90661) | 283 (90681) | 277 (90760) |
| | the median | 219 (60479) | 219 (60501) | 219 (60506) | 217 (60494) |
| | the 3rd Qu. | 148 (30253) | 145 (30250) | 141 (30254) | 144 (30254) |

**Table S1:** True causal variants retained after filtering on sample correlation, varying sample size. Uncorrelated true variants are chosen from across the genome. Number of the true SNPs retained after screening is reported (in the parenthesis is the total number of the retained variants).

| | threshold | 500 | 1000 | 2000 | 3000 |
|---|---|---|---|---|---|
| 5 snps | the 1st Qu. | 5 (90501) | 5 (90755) | 5 (90760) | 5 (90759) |
| | the median | 5 (60343) | 5 (60506) | 5 (60495) | 5 (60505) |
| | the 3rd Qu. | 3 (30239) | 3 (30245) | 3 (30254) | 3 (30254) |
| 25 snps | the 1st Qu. | 25 (89728) | 25 (90755) | 25 (90743) | 25 (90759) |
| | the median | 22 (60046) | 22 (60410) | 22 (60501) | 22 (60485) |
| | the 3rd Qu. | 12 (30253) | 13 (30254) | 13 (30254) | 12 (30254) |
| 100 snps | the 1st Qu. | 96 (90063) | 96 (90757) | 97 (90671) | 97 (90752) |
| | the median. | 86 (60489) | 88 (60458) | 94 (60507) | 90 (60505) |
| | the 3rd Qu. | 56 (30254) | 55 (30240) | 57 (30263) | 57 (30254) |
| 300 snps | the 1st Qu. | 284 (90632) | 291 (90760) | 289 (90742) | 289 (90760) |
| | the median | 233 (60497) | 254 (60497) | 268 (60506) | 264 (60507) |
| | the 3rd Qu. | 165 (30250) | 168 (30252) | 164 (30252) | 166 (60254) |

**Table S2:** True causal variants retained after filtering on sample correlation, varying sample size. The true variants are chosen from a single gene (*pbpX*). Number of the true SNPs retained after screening is reported (in the parenthesis is the total number of the retained variants).

| | threshold | 500 | 1000 | 2000 | 3000 |
|---|---|---|---|---|---|
| 50 snps (LD-prune) | the 1st Qu. | 49 (90488) | 50 (90595) | 50 (90757) | 50 (90737) |
| | the median | 43 (60450) | 45 (60438) | 45 (60479) | 41 (60491) |
| | the 3rd Qu. | 32 (30254) | 33 (30251) | 33 (30253) | 33 (30254) |
| 50 snps (gene *pbpX*) | the 1st Qu. | 49 (90589) | 49 (90753) | 49 (90759) | 49 (90756) |
| | the median | 43 (60054) | 45 (60501) | 46 (60507) | 45 (60465) |
| | the 3rd Qu. | 34 (30251) | 32 (30243) | 33 (30254) | 32 (30253) |
| 50 snps (50% gene *pbpX*, 50% gene *penA*) | the 1st Qu. | 47 (90733) | 48 (90753) | 48 (90754) | 48 (90750) |
| | the median. | 40 (60495) | 44 (60123) | 43 (60506) | 42 (60490) |
| | the 3rd Qu. | 25 (30158) | 27 (30253) | 27 (30254) | 26 (30254) |
| 50 snps (16 from gene *pbpX*, 17 from gene *penA*, 17 from gene *penX*) | the 1st Qu. | 49 (90720) | 49 (90759) | 49 (90752) | 49 (90757) |
| | the median | 46 (59328) | 47 (60459) | 46 (60471) | 46 (60499) |
| | the 3rd Qu. | 34 (30232) | 34 (30254) | 35 (30254) | 35 (30254) |

**Table S3:** True causal variants retained after filtering on sample correlation, varying sample size. The true variants are chosen from a mixture of different region types. Number of the true SNPs retained after screening is reported (in the parenthesis is the total number of the retained variants).

| | threshold | $h^2 = 0.1$ | $h^2 = 0.3$ | $h^2 = 0.6$ | $h^2 = 0.9$ |
|---|---|---|---|---|---|
| 50 snps (gene *pbpX*) | the 1st Qu. | 44 (90760) | 44 (90760) | 45 (90760) | 44 (90760) |
| | the median | 44 (60507) | 44 (60507) | 44 (60507) | 44 (60507) |
| | the 3rd Qu. | 42 (29977) | 41 (30234) | 42 (30180) | 43 (30254) |
| 50 snps (50% gene *pbpX*, 50% gene *penA*) | the 1st Qu. | 44 (90760) | 45 (90760) | 45 (90760) | 45 (90760) |
| | the median. | 43 (60507) | 43 (60507) | 43 (60494) | 43 (60507) |
| | the 3rd Qu. | 42 (30254) | 41 (30254) | 41 (30254) | 41 (30254) |
| 50 snps (16 from gene *pbpX*, 17 from gene *penA*, 17 from gene *penX*) | the 1st Qu. | 49 (90760) | 48 (90760) | 48 (90760) | 48 (60760) |
| | the median | 48 (60507) | 47 (60507) | 47 (60507) | 47 (60506) |
| | the 3rd Qu. | 41 (30254) | 42 (30253) | 42 (30254) | 43 (30254) |

**Table S4:** True causal variants retained after filtering on sample correlation, varying heritability. The true variants are chosen from a mixture of different region types. Number of the true SNPs retained after screening is reported (in the parenthesis is the total number of the retained variants).

**Table S5:** Prediction accuracy on the *Mycobacterium tuberculosis* dataset. For resistance each of the four front-line treatments we fitted a model with and without sequence reweighting and compared accuracy on a uniform random test:training split. TP – true positives; TN – true negatives; FP – false positives; FN – false negatives. The number of samples in each lineage is: 1 – 452; 2 – 448; 3 – 207; 4 – 73.

| Drug | Model | Variants Selected | Lineage | $R^2$ | TP | TN | FP | FN |
|---|---|---|---|---|---|---|---|---|
| Rifampicin | Weighted | 90 | All | 0.84 | 752 | 388 | 32 | 11 |
| | | | 1 | 0.42 | 63 | 354 | 26 | 9 |
| | | | 2 | 0.76 | 420 | 22 | 4 | 2 |
| | | | 3 | 0.83 | 201 | 5 | 1 | 0 |
| | | | 4 | 0.79 | 68 | 4 | 1 | 0 |
| | Unweighted | 132 | All | 0.86 | 755 | 388 | 29 | 8 |
| | | | 1 | 0.42 | 65 | 356 | 24 | 7 |
| | | | 2 | 0.76 | 421 | 23 | 3 | 1 |
| | | | 3 | 0.83 | 201 | 5 | 1 | 0 |
| | | | 4 | 0.79 | 68 | 4 | 1 | 0 |
| Isoniazid | Weighted | 151 | All | 0.85 | 657 | 487 | 38 | 6 |
| | | | 1 | 0.52 | 43 | 394 | 18 | 1 |
| | | | 2 | 0.69 | 389 | 59 | 14 | 5 |
| | | | 3 | 0.80 | 166 | 19 | 4 | 0 |
| | | | 4 | 0.85 | 59 | 15 | 2 | 0 |
| | Unweighted | 185 | All | 0.84 | 657 | 484 | 41 | 6 |
| | | | 1 | 0.45 | 43 | 391 | 21 | 1 |
| | | | 2 | 0.69 | 389 | 59 | 14 | 5 |
| | | | 3 | 0.80 | 166 | 19 | 4 | 0 |
| | | | 4 | 0.85 | 59 | 15 | 2 | 0 |
| Ethambutol | Weighted | 32 | All | 0.59 | 799 | 278 | 57 | 41 |
| | | | 1 | 0.13 | 89 | 272 | 39 | 41 |
| | | | 2 | 0.25 | 415 | 6 | 15 | 0 |
| | | | 3 | - | 209 | 0 | 0 | 0 |
| | | | 4 | - | 86 | 0 | 3 | 0 |
| | Unweighted | 258 | All | 0.65 | 803 | 289 | 46 | 37 |
| | | | 1 | 0.27 | 97 | 277 | 34 | 33 |
| | | | 2 | 0.25 | 411 | 10 | 11 | 4 |
| | | | 3 | - | 209 | 0 | 0 | 0 |
| | | | 4 | 0.67 | 86 | 2 | 1 | 0 |
| Pyrazinamide | Weighted | 192 | All | 0.61 | 775 | 202 | 39 | 33 |
| | | | 1 | 0.28 | 82 | 189 | 21 | 32 |
| | | | 2 | 0.34 | 413 | 10 | 14 | 1 |
| | | | 3 | - | 195 | 0 | 2 | 0 |
| | | | 4 | 0.58 | 85 | 3 | 2 | 0 |
| | Unweighted | 328 | All | 0.66 | 781 | 204 | 37 | 27 |
| | | | 1 | 0.36 | 90 | 187 | 23 | 24 |
| | | | 2 | 0.43 | 411 | 14 | 10 | 3 |
| | | | 3 | - | 195 | 0 | 2 | 0 |
| | | | 4 | 0.58 | 85 | 3 | 2 | 0 |

### 2 Supplementary figures

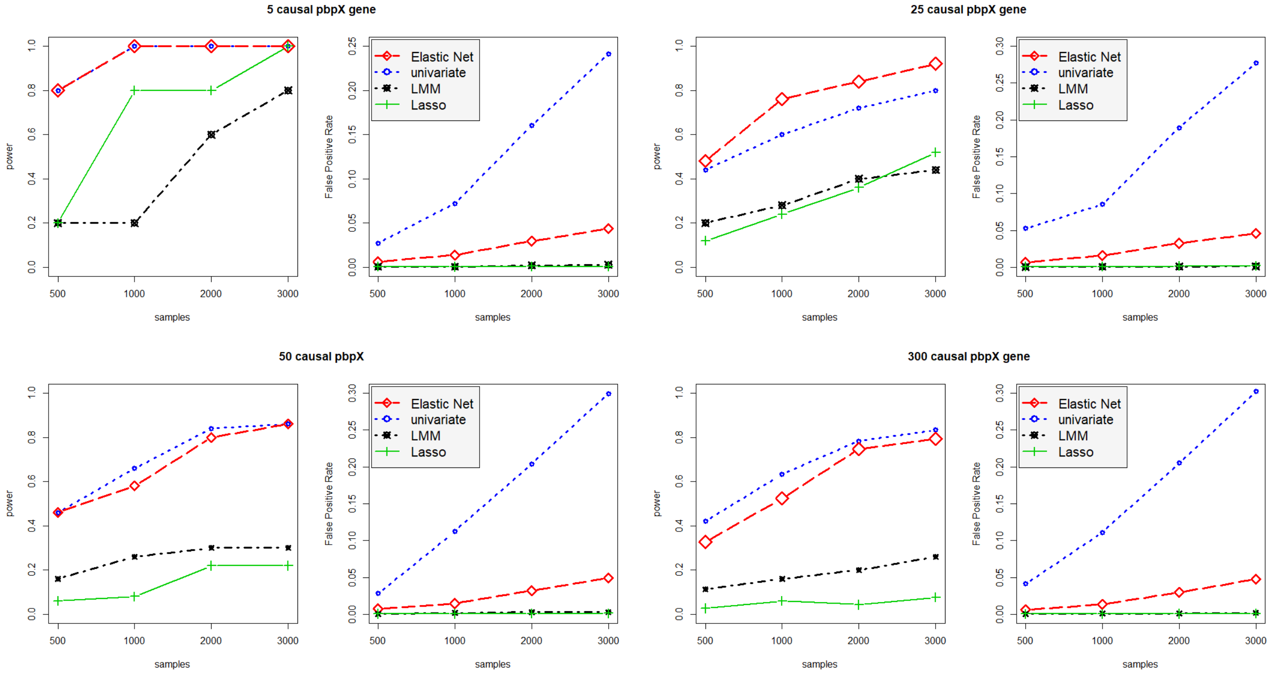


**Fig. S1:** Power and false positive rate of univariate and whole-genome regressions. Different numbers of true variants were chosen from the *pbpX* gene, sample size was varied.

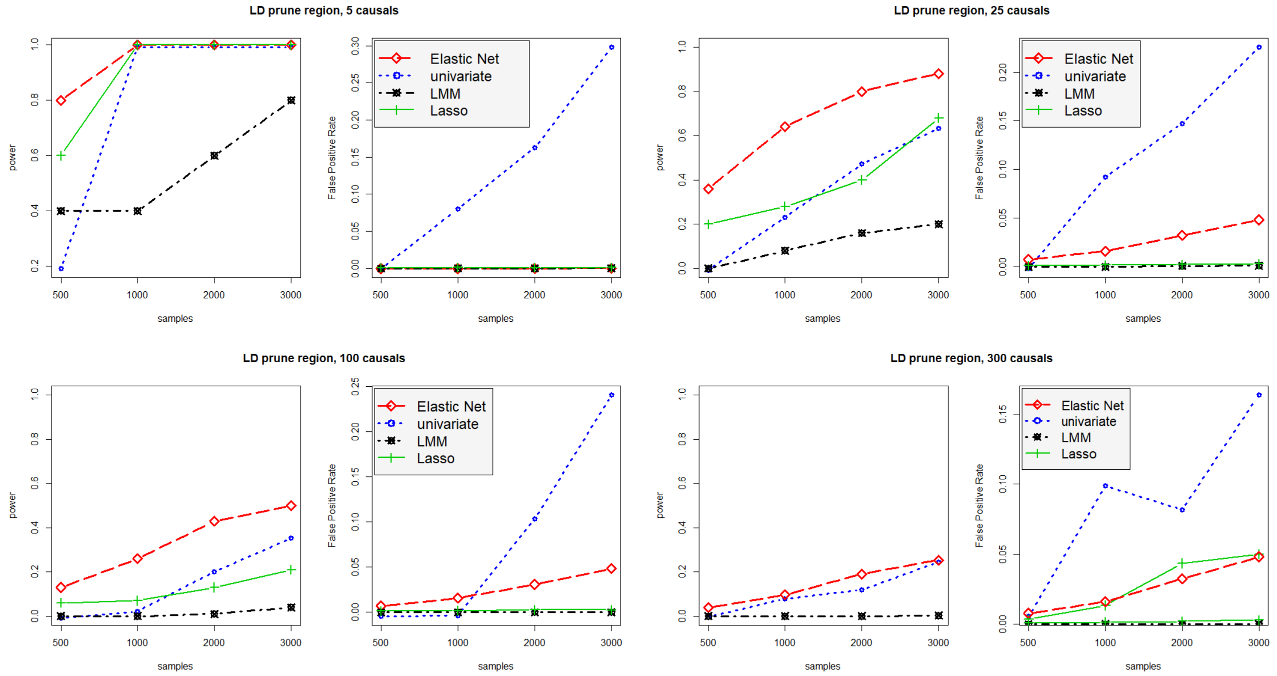


**Fig. S2:** Power and false positive rate of univariate and whole-genome regressions. Different numbers of true variants were chosen from across the genome, sample size was varied.

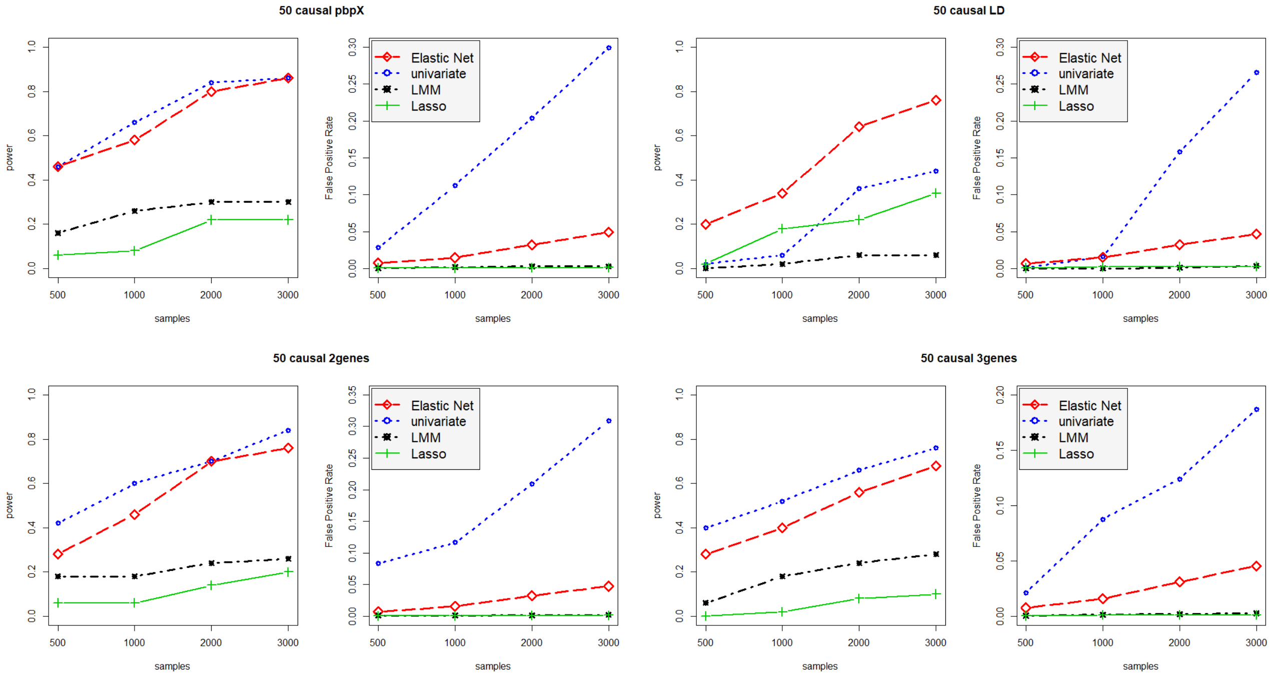


**Fig. S3:** Power and false positive rate of univariate and whole-genome regressions. 50 true variants were chosen from different set ups: LD-pruned variants across the genome, and the genes *pbpX*, *pbp1a* and *penA*. Sample size was varied.

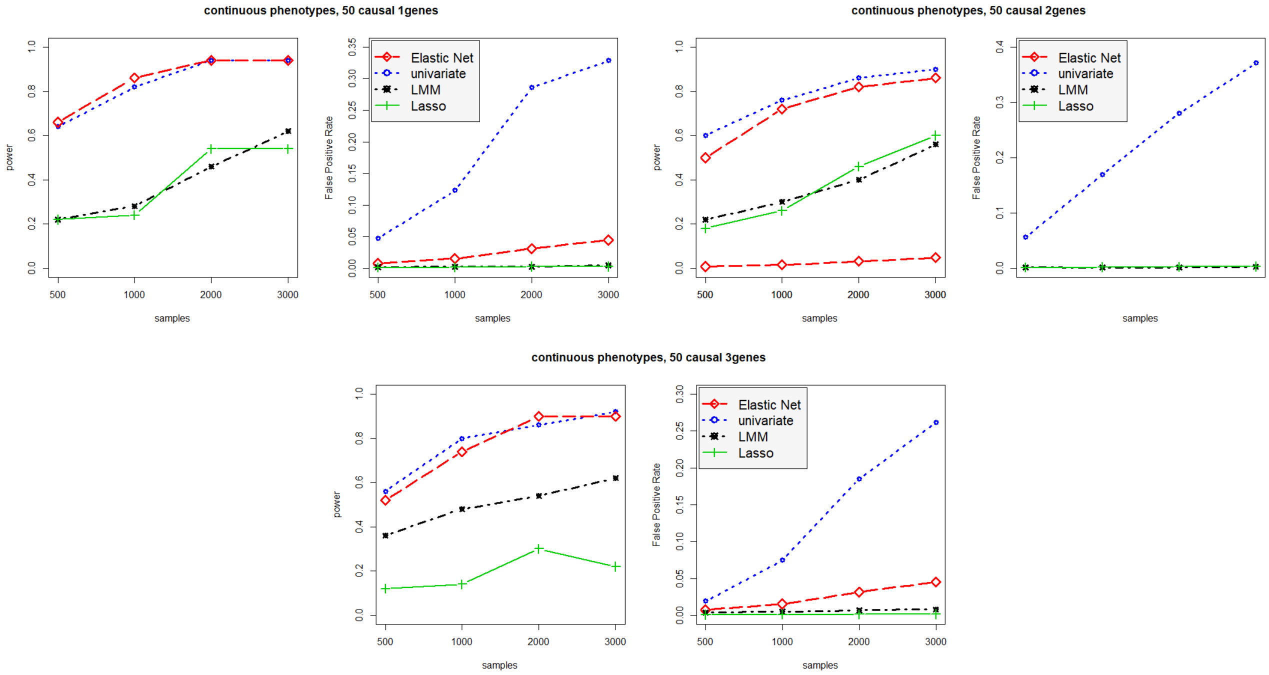


**Fig. S4:** Power and false positive rate of univariate and whole-genome regressions with a continuous phenotype. 50 true variants were chosen from different set ups: LD-pruned variants across the genome, and the genes *pbpX*, *pbp1a* and *penA*. Sample size was varied

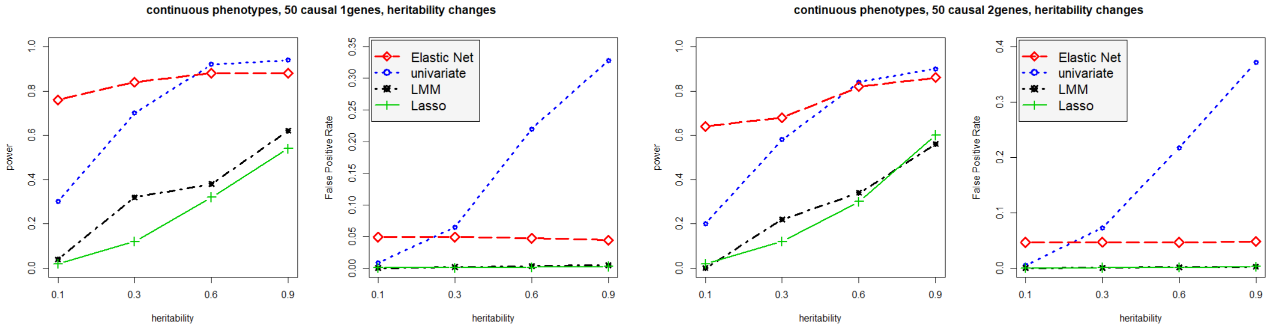


**Fig. S5:** Power and false positive rate of univariate and whole-genome regressions varying heritability of a continuous phenotype. 50 true variants were chosen from one or two genes. Sample size was fixed at 3 000.

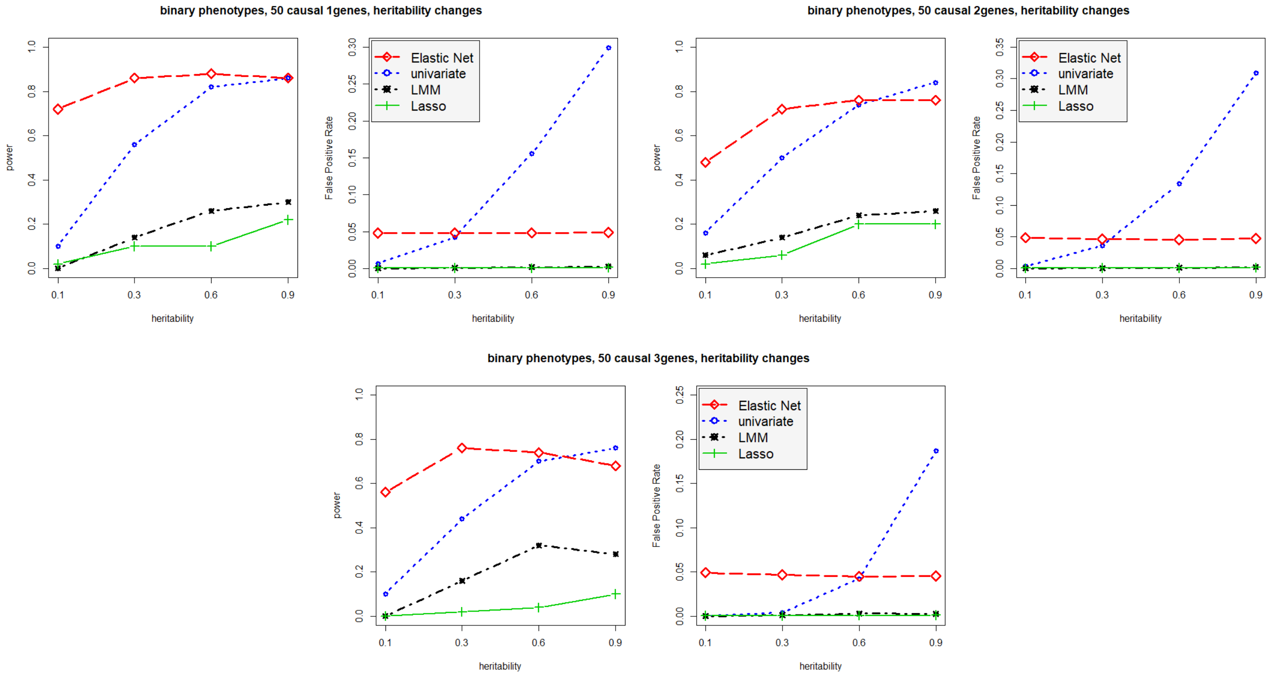


**Fig. S6:** Power and false positive rate of univariate and whole-genome regressions varying heritability of a binary phenotype. 50 true variants were chosen from one, two or three genes. Sample size was fixed at 3 000.

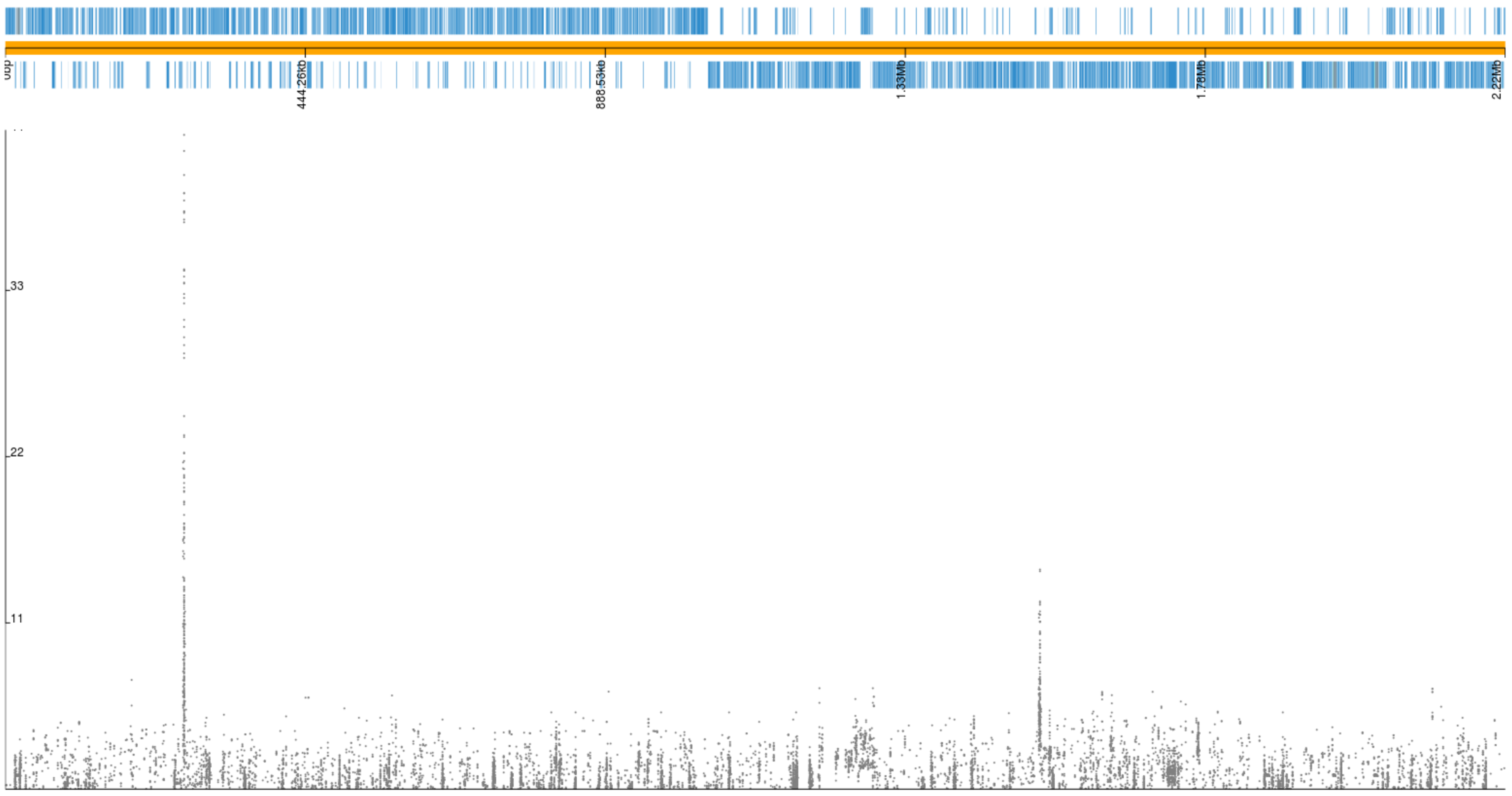


**Fig. S7:** Manhattan plot for trimethoprim resistance in *S. pneumoniae*. Plotted variants are unitigs selected by the elastic net using sequence reweighting, p-values are from the LMM.

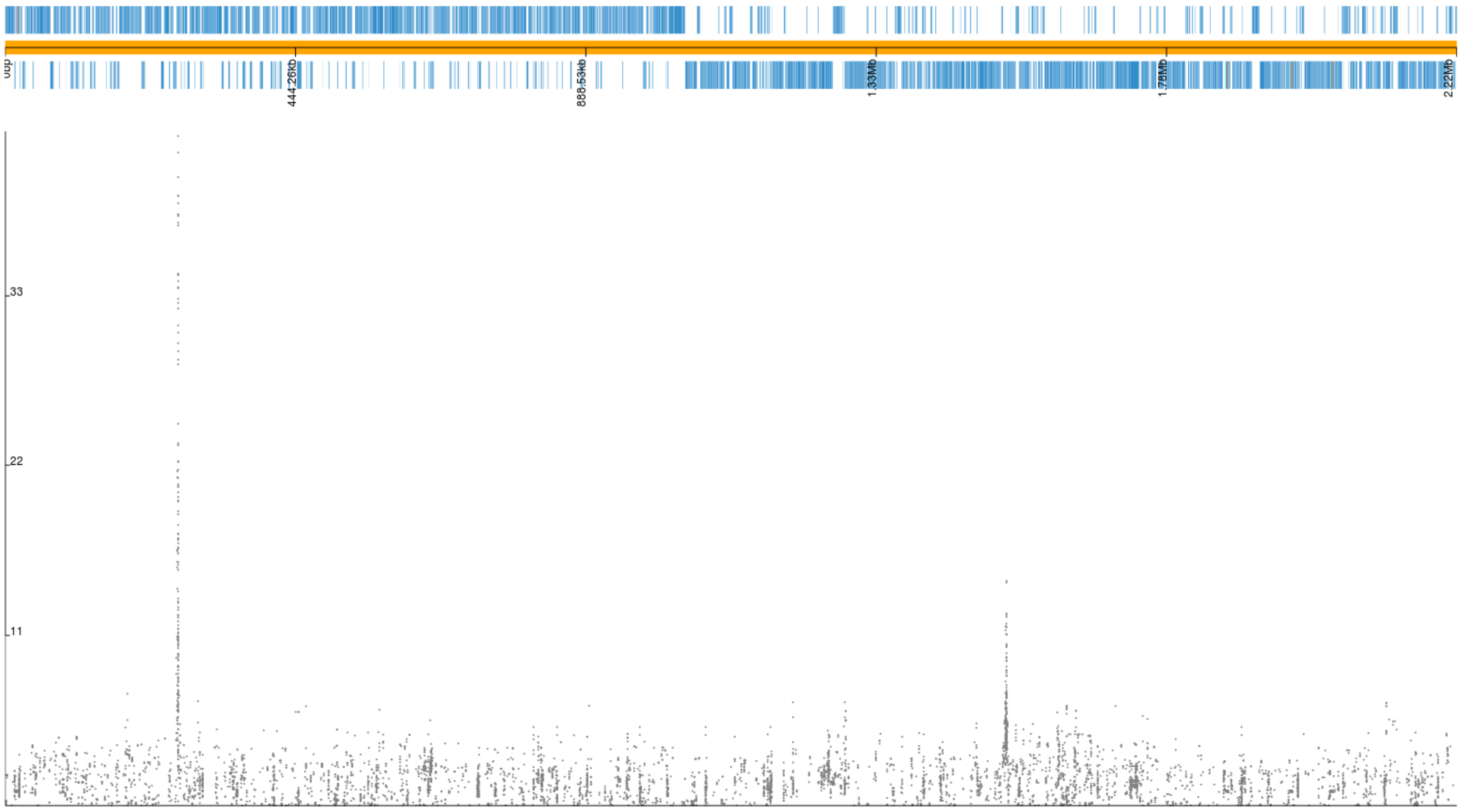


**Fig. S8:** Manhattan plot for trimethoprim resistance in *S. pneumoniae*. Plotted variants are unitigs selected by the elastic net without sequence reweighting, p-values are from the LMM.

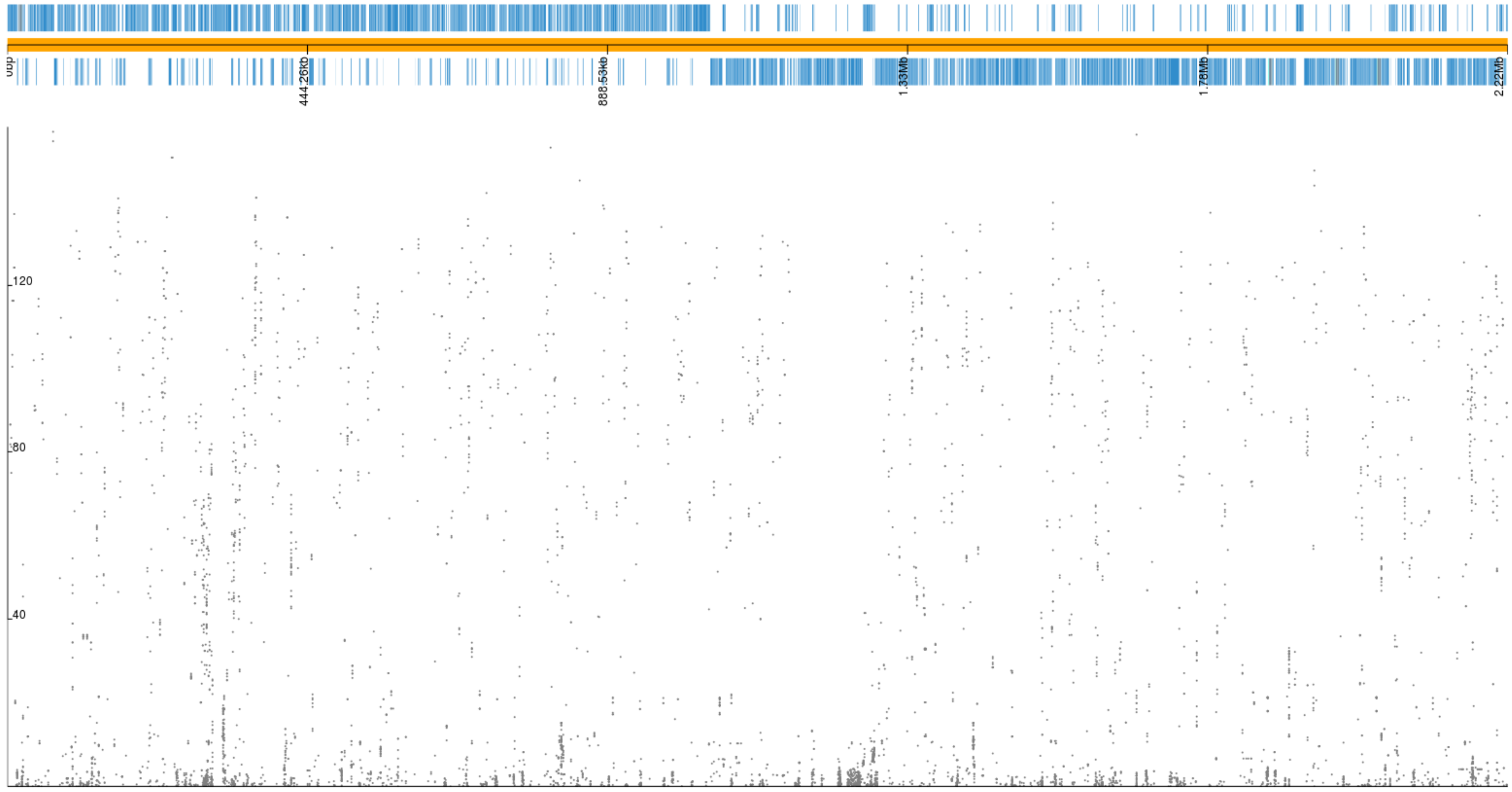


**Fig. S9:** Manhattan plot for erythromycin resistance in *S. pneumoniae*. Plotted variants are unitigs selected by the elastic net with sequence reweighting, p-values are from the LMM.

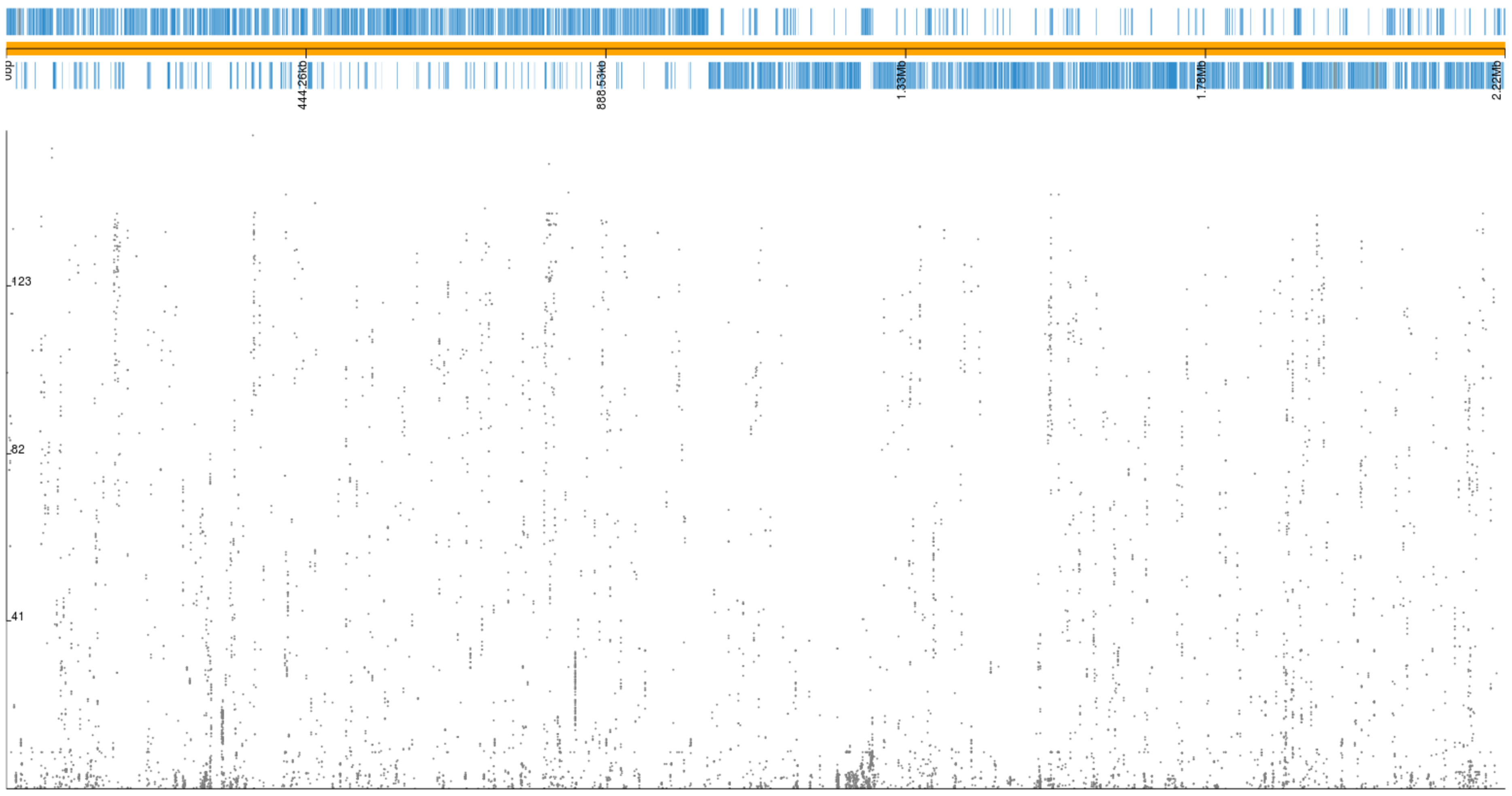


**Fig. S10:** Manhattan plot for erythromycin resistance in *S. pneumoniae*. Plotted variants are unitigs selected by the elastic net without sequence reweighting, p-values are from the LMM.

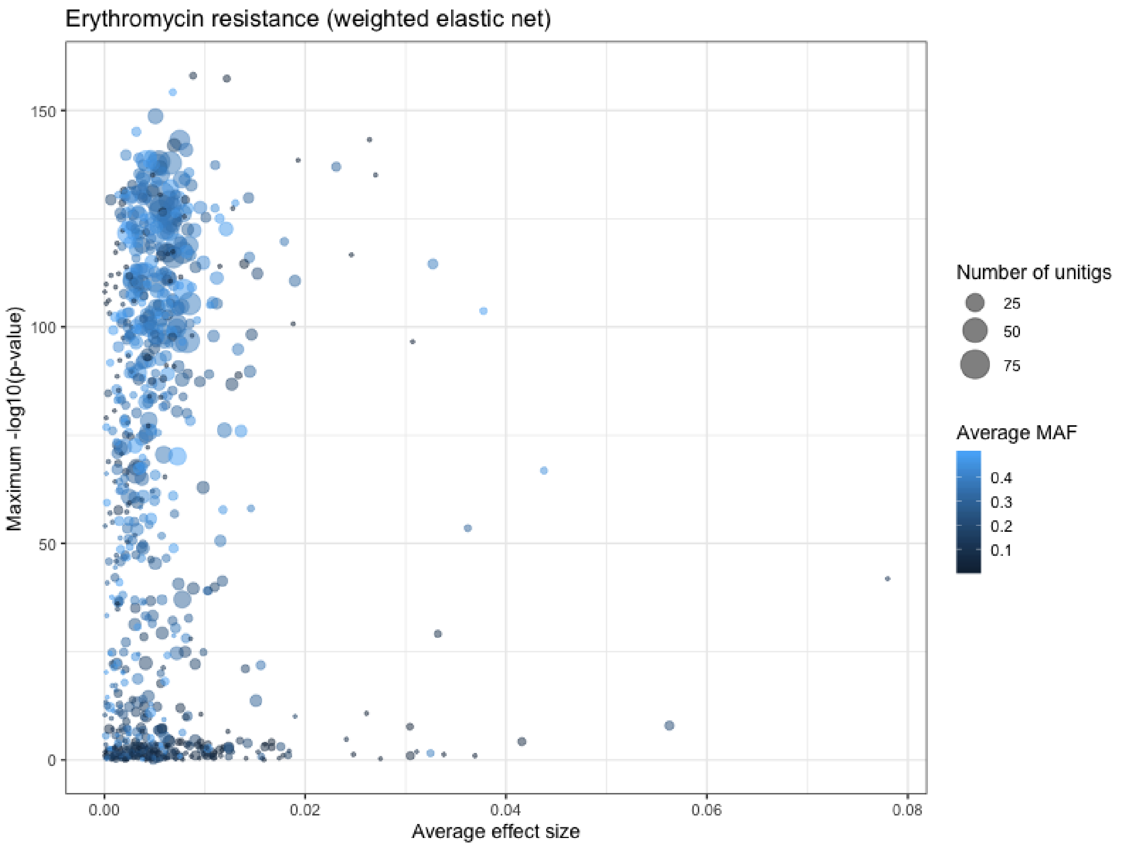


**Fig. S11:** Summary of genes with overlapping selected unitigs in the weighted GWAS. Each point is a gene, x-axis is the average effect size (beta) of unitigs covering the locus, y-axis in the minimum p-value of any unitig in the locus, size relates to the total number of unitigs mapped to the gene, colour is the average MAF of the mapped unitigs.

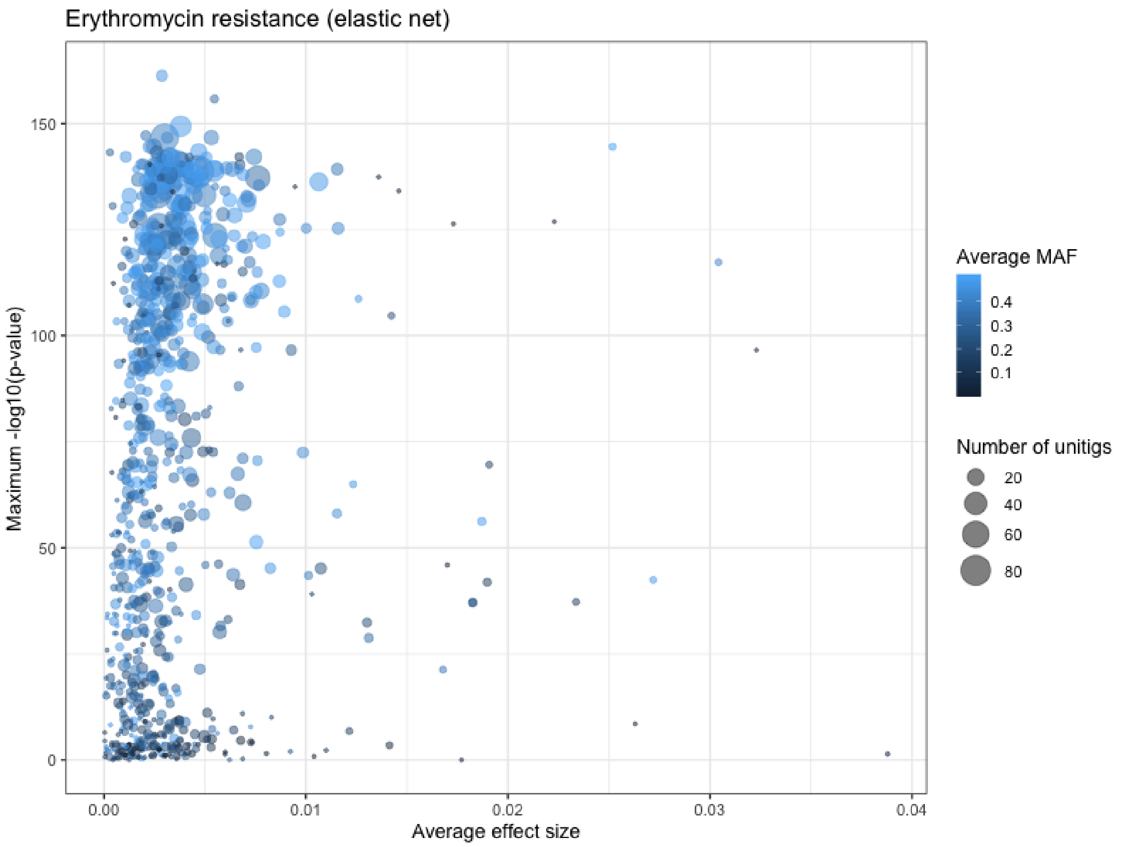


**Fig. S12:** Summary of genes with overlapping selected unitigs in the unweighted GWAS. Each point is a gene, x-axis is the average effect size (beta) of unitigs covering the locus, y-axis in the minimum p-value of any unitig in the locus, size relates to the total number of unitigs mapped to the gene, colour is the average MAF of the mapped unitigs.
